# Asynchronous excitatory neuron development in an isogenic cortical spheroid model of Down syndrome

**DOI:** 10.1101/2022.04.30.490174

**Authors:** Zhen Li, Jenny A. Klein, Sanjeev Rampam, Ronni Kurzion, Yesha Patel, Tarik F Haydar, Ella Zeldich

## Abstract

The intellectual disability (ID) in Down syndrome (DS) is thought to result from a variety of developmental deficits such as alterations in neural progenitor division, neurogenesis, gliogenesis, cortical architecture, and reduced cortical volume. However, the molecular processes underlying these neurodevelopmental changes are still elusive, preventing an understanding of the mechanistic basis of ID in DS. In this study, we used a pair of isogenic (trisomic and euploid) induced pluripotent stem cell (iPSC) lines to generate cortical spheroids (CS) that model the impact of trisomy 21 on brain development. CS contain neurons, astrocytes, and oligodendrocytes and they are widely used to approximate early neurodevelopment. Using single cell RNA sequencing (scRNA-seq), we uncovered cell type-specific transcriptomic changes in the trisomic CS. In particular, we found that excitatory neuron populations were most affected and that a specific population of cells with a transcriptomic profile resembling layer IV cortical neurons displayed the most profound divergence in developmental trajectory between trisomic and euploid genotypes. We also identified candidate genes potentially driving the developmental asynchrony between trisomic and euploid excitatory neurons. Direct comparison between the current isogenic CS scRNA-seq data and previously published datasets revealed several recurring differentially expressed genes between DS and control samples. Altogether, our study highlights the power and importance of cell type-specific analyses within a defined genetic background, coupled with broader examination of mixed samples, to comprehensively evaluate cellular phenotypes in the context of DS.

## 1 Introduction

Down syndrome (DS) is the most common genetic form of intellectual disability (ID), caused by triplication of human chromosome 21 (HSA21), with a prevalence of 1 in 700 live births in the United States (Mai et al., 2019). HSA21 contains more than 310 genes, and its triplication causes wide-spread molecular and cellular changes that underlie the characteristic phenotypes associated with DS (Vohr et al., 1989; Olmos-Serrano et al., 2016). The ID in individuals with DS is presumed to arise from anatomical and physiological alterations of the brain during atypical neurodevelopment. Histological abnormalities in brains from individuals with DS are evident as early as late-gestation, including delayed cortical lamination, reduced cerebral volume, hypocellularity, and altered neural processes (Ábrahám et al., 2012; Haydar and Reeves, 2012; Olmos-Serrano et al., 2016). These anatomical changes are, in turn, a product of changes in the embryonic brain, including abnormal divisions of neural progenitors, aberrant neuronal migration, and altered cell-to-cell adhesion(Tyler and Haydar, 2013; Huo et al., 2018; Bells et al., 2019). However, molecular processes underlying these cellular, anatomical and physiological changes that result in ID have not been fully elucidated yet.

On one hand, the lack of mechanistic knowledge is due in part to the limited access to and ethical considerations of conducting research in human brain tissue, which restricts our ability to temporally examine how trisomy affects the development of different types of brain cells. On the other hand, mouse models of DS, while invaluable, are challenged by inconsistency in genetic backgrounds, reduced mutation penetration, and phenotypic drift(Gardiner et al., 2003; Gupta et al., 2016; Kazuki et al., 2020; Shaw et al., 2020). Thus, to model human- and disease-relevant aspects of DS, *in vitro* cultures of human induced pluripotent stem cells (iPSCs) have risen in popularity, due to their ability to reflect regional and cell type-specific features of the human brain. Three dimensional (3D) cortical spheroids (CS) have been shown to surpass two dimensional (2D) iPSC cultures in recapitulating signaling pathways, patterning, fate acquisition, and developmental trajectories of the *in vivo* environment (Kathuria et al., 2020). CS have also been shown to better preserve the expression of cell adhesion molecules, extracellular matrix components, and cell membrane structures (Scuderi et al., 2021) and possess a greater transcriptomic overlap with human fetal brain at mid-term gestation (Pasca et al., 2015; Qian et al., 2019; Kathuria et al., 2020).

In this study, we used a pair of isogenic (euploid and trisomic) iPSCs derived from an adult female with DS to generate iPSC-derived CS, following a recently published protocol (Madhavan et al., 2018). In addition to morphological and histological examination, we also performed single cell RNA sequencing (scRNA-seq) to characterize molecular alterations at the single cell level of resolution. While our CS contained seven major cell types, including radial glial cells (RGCs), intermediate precursors (IPCs), astrocytes, and inhibitory neurons, our transcriptomic analysis identified excitatory neuron (ExN) clusters as the most affected by trisomy. Specifically, our studies identified a cluster of cells corresponding transcriptionally to layer IV cortical neurons (ExN4) as the major dysregulated cell type affected by trisomy 21. ExN4 displayed profound developmental divergence from the corresponding euploid cluster, including many differentially expressed (DEX) genes and affected processes related to neuronal motility and establishment of cortical architecture. The dataset also revealed gene candidates in specific cell types that drive alterations in developmental trajectories.

We then performed a direct comparison of our scRNA-seq study to previous datasets generated from the same isogenic lines as well as from human postmortem brain tissue (Olmos-Serrano et al., 2016; Palmer et al., 2021; Nava et al., 2022). This comparison revealed that despite differences in technical approaches and the source of trisomic samples, there is a portion of shared HSA21 and non-HSA21 genes affected in all the studies. This analysis also identified transcriptomic divergence and distinct transcriptional profiles relating to the specific genetic background of the individual (sex, allelic composition). By comparing the current CS dataset to previously published studies, we demonstrate the benefit of using isogenic cell lines in uncovering consistent biological factors across studies and platforms.

## 2. Materials and Methods

### 2.1 Generation of CS

We received a pair of isogenic lines, consisting of a trisomic line (WC-24-02-DS-M) and a euploid control (WC-24-02-DS-B), as a generous gift from Anita Bhattacharyya’s lab at the University of Wisconsin, Madison. These lines were validated previously and deposited at WiCell^®^ Research Institute (Madison, WI). IPSCs were passaged and cultured on Matrigel^®^ (Corning) using mTeSR™ plus media (StemCell Technologies^®^). Cells below passage 30 were used to generate CS. About 1.5 × 10^6^ trisomic and euploid iPSCss dissociated with accutase (StemCell Technologies) were used to generate around 100 spheroids that were differentiated further into CS following a published protocol with modifications (Madhavan et al., 2018). Briefly, the dissociated cells were transferred to individual low-adherence V-bottom 96-well plates (S-Bio Prime) in 150μl TeSR5/6 media (StemCell Technologies^®^) with 50μM Rock inhibitor Y-27632 (Tocris BioScience), 5 μM Dorsopmorphin (Tocris BioScience) and 10 μM SB-431542 (Tocris BioScience). The same media without Rock inhibitor was used and changed daily for the next five days. On day six, the media was changed to spheroid media containing Neurobasal-A media supplemented with B-27 without vitamin A (Invitrogen), Glutamax (Invitrogen), and Penicillin/Streptomycin. Basic fibroblast growth factor (FGF-2, 20ng/ml, R&D systems) and epidermal growth factor EGF (10ng/ml, R&D systems) were added to the media on days 7-24. On day 25, spheroids were transferred to ultra-low attachment 24-well plates (Corning) and 1% Geltrex (Invitrogen) was added to the media. Brain Derived Neurotrophic Factor (BDNF, 20ng/ml, R&D systems) and Neurotrophin-3 (NT-3, 20ng/ml, R&D systems) were used for neural differentiation between days 27 and 41. To expand the existing small population of oligodendrocytes in the spheroids, beginning on day 50, 10 ng/ml platelet-derived growth factor-AA (PDGF-AA, R&D systems) and insulin-like growth factor-1 (IGF-1, R&D systems) were supplemented to the media changes for 10 days. Between days 50 and 60, the media was supplemented with 40 ng/ml 3,3’,5-triiodothronine (T3, R&D systems). The spheroids were maintained in spheroid media from day 70 until completion of the experiment with half-media changes every other day. Multiple, temporally overlapping spheroid cultures were generated to provide a constant source for sampling and analysis of developmental markers. The mycoplasma contamination test was performed regularly using PCR Mycoplasma Test Kit I/C (PromoCell®).

### 2.2 Single cell dissociation and capture

CS dissociation was performed on day 130 as described. Four CS generated in different wells were pooled per sample and dissociated with Worthington Papain dissociation system (Worthington Biochemical Corp., Lakewood NJ, Cat#: LK003150) following the protocol by the manufacturer. Prior to dissociation, we oxygenated the papain solution with 95% O2 and 5% CO2 to insure cell viability. The CS were first cut into small pieces and then dissociated in 20 units/ml papain and 0.005% DNase solution at 37°C with thorough constant agitation for 40 minutes. The mixture was titrated with 5 ml pipette and the cell suspension was centrifuged at 300g for 2 minutes at room temperature. The pellet was resuspended with PBS containing 1% BSA. Cell viability and number was assessed using Tripan-Blue on Countess automatic cell counter (Invitrogen). Cell samples at a concentration of 1000 cells/ul were submitted for a single cell capture. 10X Genomics Chromium^®^ single cell preparation system was used for cell capture following manufacturer’s protocol.

### 2.3 Library preparation and sequencing

The synthesis of cDNA, cDNA amplification, and the preparation of the libraries were performed using the 10x Genomics Chromium Single Cell 3’ Library and Bead Kit (v3). according to manufacturer’s instruction. Sequencing was done on NovaSeq 6000 at the Single Cell Sequencing Core at Boston University School of Medicine.

### 2.4 Read alignment

Fastq files containing pair-end reads of each sample were aligned to GRCh38 Genome Reference Consortium Human Reference 38 (hg38) and GENCODE annotation (v35) using Cellranger (v3.1.0) *count* function with default settings. Cellranger *aggr* function was then used to combine aligned and filtered count matrix from all samples.

### 2.5 Bioinformatics analyses

#### 2.5.1 Quality control

Cells with 1) number of detected genes greater than 1000 and 2) percentage of reads mapped to mitochondrial genome between 1% to 10% were kept. After filtering cells, only protein coding genes in each cell were used for downstream analyses. Mitochondrial genes were removed. Data were normalized using *NormalizeData* function from the *Seurat* R software package with normalization method set to “LogNormalize” and scale factor as 10,000 (Stuart et al., 2019).

#### 2.5.2 Dimension reduction and clustering

To perform dimenstion reduction and clustering, we first identified the top 2000 highly variable genes (HVGs) using *FindVariableGenes* function from the *Seurat* R software package. The HVGs were scaled before being applied to principal component analysis (PCA) as input. Top 10 principal components (PCs) with the highest standard deviation were used to perform UMAP dimension reduction resulting in a 2D representation of the dataset. Clustering was done first by calculating the neighborhood of each cell with *FindNeighbors* function on the two UMAP coordinates with k parameter set to 15. Then, *FindClusters* function was called with resolution set to 0.15.

#### 2.5.3 Differential expression (DEX) analyses

We conducted DEX analyses using Seurat function *FindAllMarkers*. We took cells from one cell type and compared it to the rest of all the cells, using a binomial model. For any given comparison, we only considered genes that were expressed by at least 25% of cells in either population. Genes that exhibit adjusted p-values under 0.1 were considered statistically significant. The Database for Annotation, Visualization, and Integrated Discovery (DAVID) v6.8 was used for gene ontology (GO) analysis. Briefly, all statistically significant genes for each cell cluster were entered into the database and statistically significant biological processes associated with the gene lists were identified (FDR < 0.05). Biological processes were reported in order of fold enrichment, or the ratio of the DEX genes in the list involved in a particular process to the total number of genes that could be involved in that process in *Homo sapiens*.

#### 2.5.4 Diffusion map

To generate diffusion map (DM) for all cells in the dataset, we first selected the top 500 HVGs and performed PCA as described in the previous section. The top 20 PCs were used to find the optimal sigma (σ) using function *find_sigmas* from R package Destiny with default parameters (Angerer et al., 2016). Then, the top 20 PCs were used as input in function *DiffusionMap*, with 2σ as the diffusion scale parameter and number of nearest neighbors (k) set as 100. To calculate DM for each individual cell type, the abovementioned procedure was followed with data within each cell type as input and k set to 25.

#### 2.5.5 Pseudotime analysis

We used R package URD following recommended steps with minor adjustments based on the structure of the dataset (Farrell et al., 2018). Briefly, a subset of aRGC1 cells near the center of the DM were set as the root. The DM was flooded 100 times to establish the pseudotime axis. Tips of the DM were identified from the final stage of pseudodevelopment. Biased random walks were then performed from each tip for 10000 times. Lastly, a tree graph was built using *buildTree* function with default settings except threshold of *p*-value set to 0.05.

#### 2.5.6 Principal graph analysis

To identify genes associated with different regions of DM, we first manually converted our dataset from a URD object into a monocle object (Cao et al., 2019). Then, the function *graph_test* from R package monocle3 was performed on the monocle object. Moran’s I greater than 0.3 and adjusted *p*-value less than 0.01 were used as threshold to identify genes associated with either trisomic or euploid cells on the DM.

#### 2.5.7 Inter-genotype distance

To assess the genotypic differences in each cell type, we first calculated the Euclidean distances between each trisomic cell to each euploid cell within each cell type using *dist* function on DM, which were then averaged to get observed inter-genotype distance (oIGD). We then performed 1000 permutations within each cell type. During each permutation, the genotype labels were randomized within each cell type, and an inter-genotype distance (eIGD) was calculated by the same process as oIGD. Lastly, a *p*-value was calculated for oIGD based on the distribution of eIGDs of the same cell type. To compare between cell types, oIGD and eIGDs of each cell type were normalized by dividing the average Euclidean distance between each unique pairs of euploid cells within the respective cell type.

### 2.6 Immunohistochemistry

For immunohistochemistry (IHC), the CS were fixed overnight with 4% ice-cold paraformaldehyde, washed three times, ten minutes each, with PBS, and cryoprotected in 30% sucrose overnight. The spheroids were embedded in 30% sucrose/Optimal Cutting Temperature compound (OCT; Sakura) at 1:1 ration and sectioned at 12μm. Sections were washed three times with PBS, blocked for 30 minutes in PBS containing 0.1% Triton X-100 (PBST) and then incubated in a blocking solution containing 5% donkey serum in PBST for an hour at room temperature. Next, the sections were incubated with the primary antibodies diluted in the blocking buffer at 4°C overnight. The next day, the slides were washed three times with PBST for 10 minutes each, followed by incubation with secondary antibody for an hour at room temperature. Then, the slides were washed three times with PBS for 10 minutes and coversliped with ProLong™ Gold Antifade Mountant with DAPI (ThermoFisher).

The following primary antibodies were used: rabbit anti-CC3 (1:750, Cell Signaling cat. number: 9661-s); mouse anti-SATB2 (1:250, Abcam, cat. number: ab51520); rat anti-CTIP2 (1:400, Abcam cat. number: ab18465); rabbit anti-FOXG1 (1:250, Abcam cat. number: ab196868); goat anti-SOX2 (1:250 R&D Systems, cat. number: AF2018), rabbit anti-TBR1 (1:250, Abcam, cat. number: Ab31940). All secondary antibodies were AlexaFluor conjugated, used at a dilution of 1:500 and obtained from LifeTechnologies.

### 2.7 Confocal Microscopy, Imaging and Quantification

For each organoid, three to four regions of cut sections were imaged per spheroid using a Zeiss LSM 710 confocal microscope system (Carl Zeiss, GER) and z-stacks (1024 × 1024 pixels) were collected using a 20x or 40x objective lens. For markers of developing neurons (SATB2, CTIP2, TBR1), the regions imaged were located in the vicinity of the ventricular-like zones present in the spheroid sections that were identified by morphology and presence of positive cells. For the other markers, regions were randomly chosen along the spheroid edge. The optical density of antibody labeling was assessed and quantified using particle analyses function through the imageJ/FIJI (RRID:SCR_003070; Medalla et al., 2017). For the analyses, the threshold for the signal was set in the first field and subsequently applied to the rest of the fields of the same image. The percent of antibody-recognized area was calculated out of the total area covered by DAPI for each field. To analyze cortical layer markers (SATB2, CTIP2, and TBR1 etc.), labeled cells in each z-stack were counted using the ACEq application, a ‘3-dimensional version’ of the app that was designed to quantitatively assess markers across the z-stack and correct for overlap as described previously (Klein et al., 2022). This version can be publicly accessed through Zeldich lab website^1^. The counts were first averaged for each region of the sliced organoid and then the values were averaged for a value of the total number of the organoids per condition to calculate a representation of mean ± standard error of the mean.

### 2.8 Statistical analyses and data presentation

For IHC experiments quantification, Graphpad Prism software was used for the plotting of the data and assessing statistical significance between the conditions. We used an unpaired two-tailed student’s T-test to compare the quantification of cortical layer markers in isogenic euploid and trisomic CS following IHC. For the measurements of the size of the organoids across different time points, one-way ANOVA with post hoc Tukey’s test was used.

Exact hypergeometric probability test was used to calculate the statistical significance of overlap between DEX genes of different datasets. Kolmogorov-Smirnov test was used to assess the distribution of cell density along pseudotime. All graphs related to bioinformatics analyses were generated with *ggplot2* R package except when noted otherwise (Wickham, 2009).

## 3 Results

### 3.1 Generation of euploid and trisomic CS containing diverse cell lineages

Following a recently published protocol (Madhavan et al., 2018), we generated dorsal forebrain fated CS from an isogenic pair of euploid (WC-24-02-DS-B) and trisomic (WC-24-02-DS-M) iPSCs derived from an adult female with DS that contained progenitors, neurons, astrocytes, and oligodendrocytes. Following continuous culturing and developmental characterizations up until 130 days, CS were transcriptomically profiled by scRNA-seq and subjected to immunohistochemistry (IHC) (**Figure 1**). We observed expression of the ectoderm and neural stem cell marker sry-box transcription factor 2 (SOX2) on day 30 in both euploid and trisomic CS enriched within rosette structures reminiscent of cortical ventricular zones (VZ) (**Figure 2B**,**C)**. In addition, we also confirmed the forebrain identity of the isogenic CS by staining with forkhead box G1 (FOXG1) (**Figure 2D**). Following continuous culturing to allow neuronal and glial differentiation, we successfully verified the presence of neurons by *microtubule associated protein 2* (MAP2) positivity and astrocytes by glial fibrillary acidic protein (GFAP) positivity starting from Day 50. Beginning on Day 80, we also observed emergence of oligodendrocytes confirmed by oligodendrocyte transcription factor 2 (OLIG2) IHC. These cell types continued to mature and were present at day 130 (**Figure 2E**).

**Figure 1.**
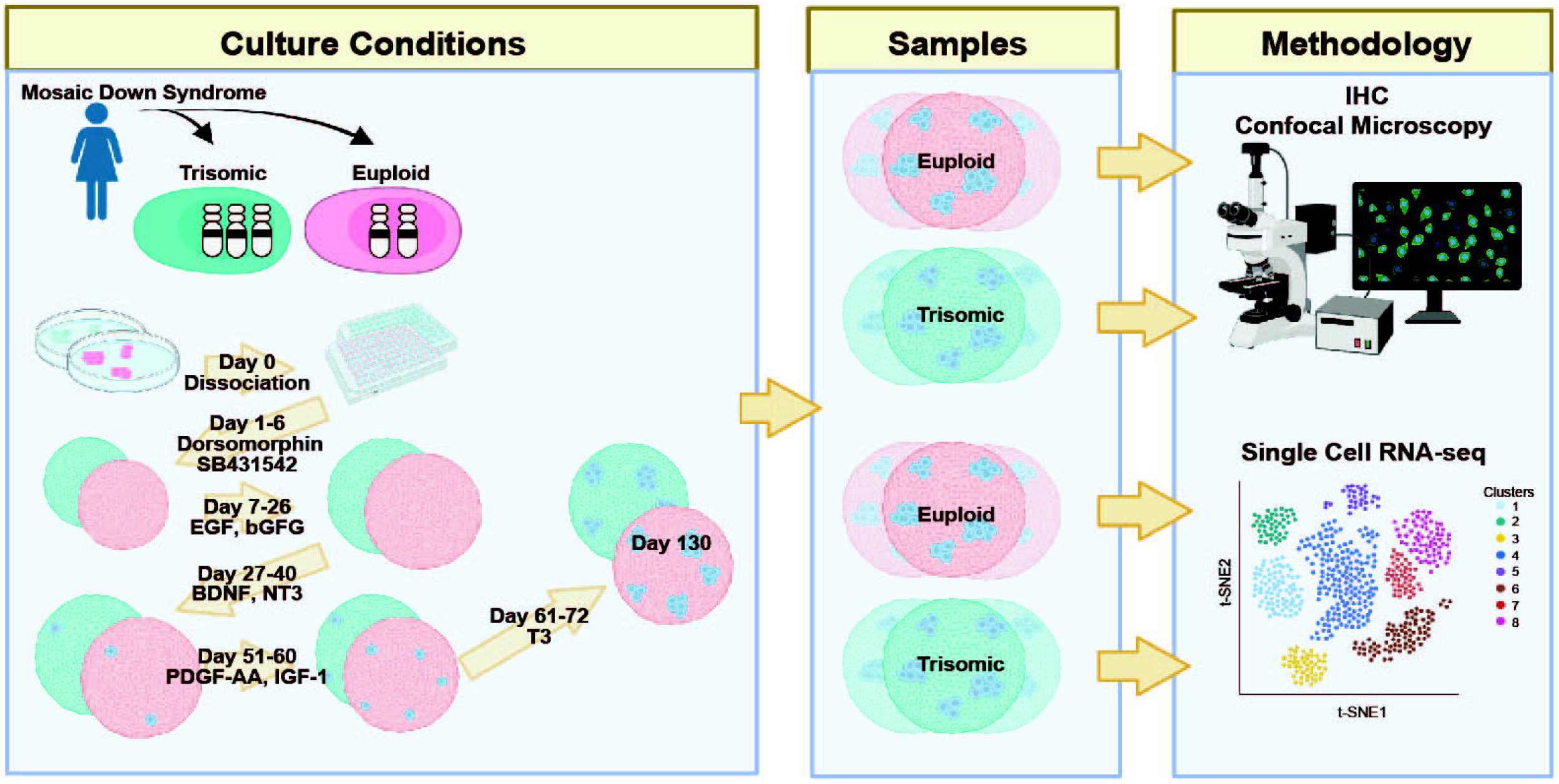
Schematic representation of the experimental protocol and study design. Isogenic HSA21 euploid (red) and trisomic (blue) induced pluripotent stem cell (iPSC) lines derived from awoman with Down Syndrome were differentiated into cortical spheroids (CS) and analyzed at day 130 via IHC and scRNA-seq. Created with BioRender.com.

**Figure 2.**
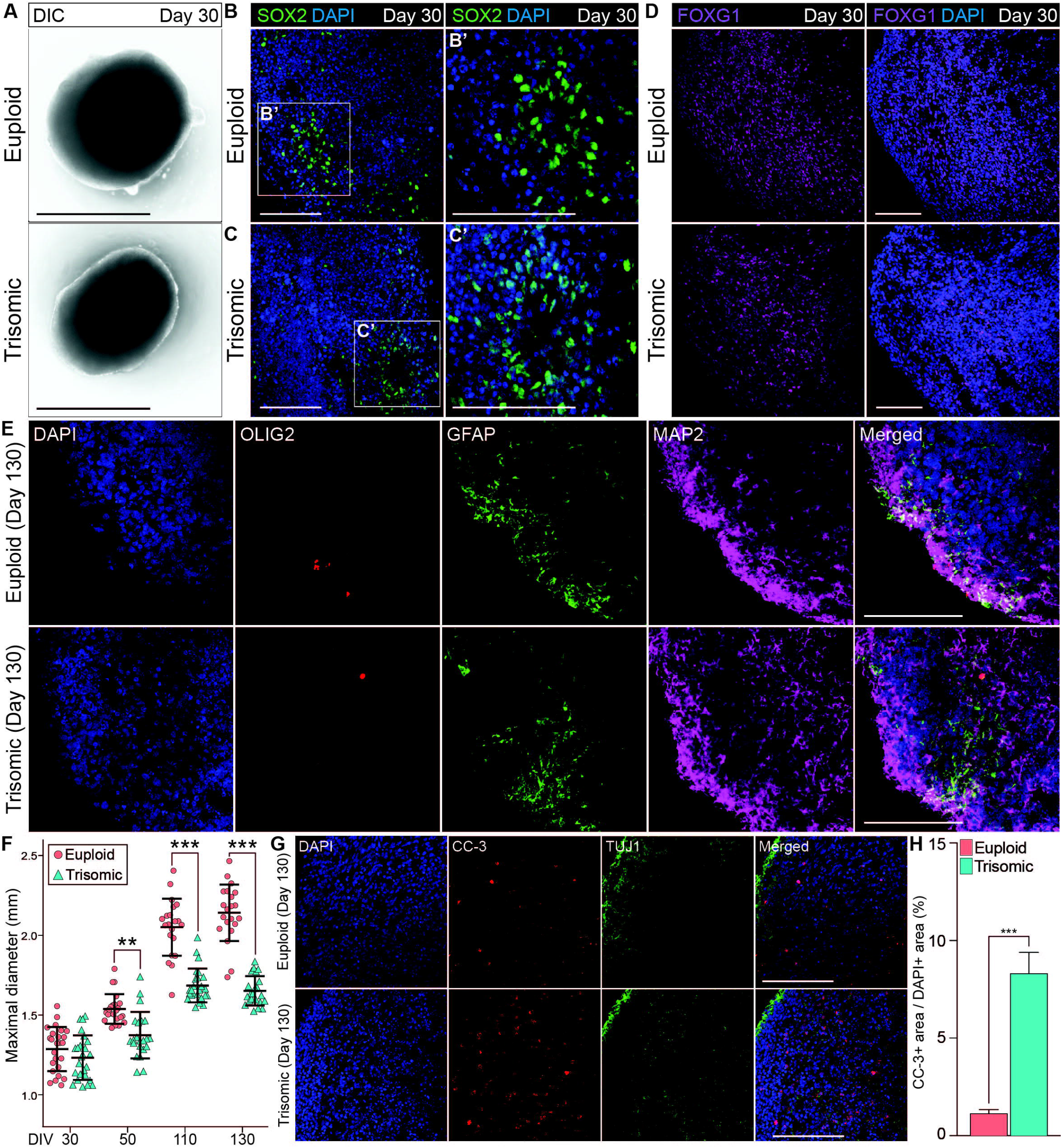
Generation and characterization of isogenic CS. **(A)** Bright field images of CS on day 30. (B, B’) IHC staining with the anit-SOX2 showing the presence of the rosette-like structures in euploid CS on day 30. **(C, C’)** IHC staining with the anit-SOX2 showing the presence of the rosette-like structures in trisomic CS on day 30. **(D)** IHC staining with FOXG1antibody on day 30 in euploid and trisomic CS. **(E)** IHC staining with anti-GFAP, anti-MAP2 and anti-OLIG2 antibodies in euploid and trisomic CS on day 130. **(F)** Jitter plot showing the distribution of CS diameters on days 30 (euploid, n = 25; trisomic, n = 25), 50 (euploid, n =23; trisomic, n = 22), 110 (euploid, n = 21; trisomic, n = 25), and 130 (euploid, n = 21; trisomic, n = 22). Euploid samples are represented by red circles, while trisomic samples are represented by blue triangles. The results are analyzed by one way ANOVA followed by Tukey’s multiple comparisons test. **(G)** IHC staining with anti-of CC-3 and anti-TUJ1 antibodies in euploid and trisomic CS on day 130. **(H)** Bar graph showing the percentage of area with CC-3 IHC signal over the area with DAPI signal quantified through particles analysis via ImageJ s and analyzed using student t-test; (euploid, n = 14; trisomic, n = 13). Error bar represents standard error. *p<0.05; **p<0.01; ***p<0.001. The quantification results are generated from three independent differentiation experiments. Scale bar is 1000 μm in A and 100 μm in B, C, D, E and G.

### 3.2 Trisomic CS display a reduced cortical volume

The euploid and trisomic CS developed in a comparable manner as measured by spheroid diameter during early stages of differentiation (day 30), preceding the induction of cortical expansion by the application of the neurotrophins BDNF and NT3 (1231μm±19.8, trisomic; 1285 μm±19.9, euploid; p=0.45). The size of the spheroids diverged upon the completion of the neurotrophin treatment at Day 50, as the diameter of trisomic CS were significantly smaller compared to euploid controls (1372μm±20.9μm, trisomic; 1536μm±13.2μm, euploid; p=0.002, **Figure 2F**). The size differences became even more pronounced on day 110, when trisomic organoids measured at an average of1684μm±15.1μm, while euploid organoids measured 2050μm±25.6μm (p<0.001, **Figure 2F**). On Day 130, trisomic organoids measured at an average of 1650μm±13.1μm, while euploid organoids at 2140μm±25.3 μm (p<0.001, **Figure 2F**). The difference in size is consistent with smaller size of embryonic bodies and brain organoids reported by other groups (Tang et al., 2021) and is in line with reduced cortical volume in individuals with DS (Wisniewski, 1990; Baburamani et al., 2020).

We hypothesized that apoptosis may be an underlying cause of the consistent decrease in size of the trisomic CS. Therefore, we first examined markers of apoptosis using IHC of cleaved caspase-3 (CC-3). We found increased CC-3 IHC signal in trisomic CS at day 90 with 10.3%±2.1% of cells in showing CC-3 positivity, whereas only 4.7±0.9% in euploid CS were positive of CC-3 (p<0.038;, **Supplemental Figure 1A,B**). On Day 130, 8.4%±1.1% cells in trisomic CS were positive for CC-3, compared to the euploid CS where only 1.2%±0.2% cells were positive (p<0.001; **Figure 2G,H**). These results suggest that the reduced size of the trisomic CS can be attributed at least in part to increased cell death.

### 3.3 ScRNA-seq analysis unravels alteration in neural development in trisomy 21 CS

To further characterize the CS, we performed scRNA-seq analysis on day 130 (**Figure 1)**. Two samples were collected for each genotype and each sample consisted of a pool of four CS from the same differentiation experiment, resulting in the processing of eight spheroids per genotype total. The samples were processed following 10X Genomics scRNA-seq protocol and an estimated number of 8890 cells were captured. After quality control (QC), 6093 cells were kept for downstream analysis. To confirm the reproducibility of sample preparation and sequencing, we compiled reads by sample and compared the genomic coverage across samples in five million base pair windows across the entire genome. We observed identical patterns of genomic coverage across the four samples, except on HSA21 where reads from the trisomic CS displayed an elevated level of disturbance compared to euploid samples (**Supplemental Figure 2**). To further confirm the effect of trisomy at the level of individual samples, we performed differential gene expression (DEX) analysis between trisomic and euploid samples using *DESeq2* program (Love et al., 2014). As expected, in trisomic samples we observed a much greater number of upregulated than downregulated genes on HSA21. In contrast, the numbers of up- or downregulated genes on the other autosomes were comparable (**Supplemental Figure 2**). At the level of individual cells, we detected an average of over 3000 genes in each cell with an average read depth (UMI) around 10000 (**Supplemental Figure 3A**). All cells that passed QC had no more than 10% and no less than 1% of total reads mapped to mitochondrial genome (**Supplemental Figure 3A**). We then performed dimension reduction and depicted the transcriptome from each cell in 2D space using UMAP (Becht et al., 2018). No batch effect or overt differences between trisomic and euploid samples were observed (**Supplemental Figure 3B**,**C**).

Next, we performed unsupervised clustering following *Seurat* v3 pipeline (Stuart et al., 2019) and identified 16 clusters representing seven major cell types (**Figure 3A**). The major cell types include apical radial glia cells (aRGC), basal radial glial cells (bRGC), intermediate progenitor cells (IPC), astrocytes (Ast), oligodendrocytes (Olig), inhibitory neurons (InN), and excitatory neurons (ExN). All cell types were present in each of the four samples (**Figure 3B**). Each cell type expressed canonical marker genes including *SOX2* (RGCs), eomesodermin (*EOMES*) (IPCs), cut like homeobox (*CUX2*, layer II/III ExN), special AT-rich sequence-binding protein (*SATB2*, layer II-IV ExN), RAR related orphan receptor B (*RORB*, layer IV ExN), BAF chromatin remodeling complex subunit (*BCL11B*, layer V ExN), glutamate decarboxylase 2 (*GAD2*, InN), *OLIG1* (Olig), as well as astrocytic markers, *GFAP* and aldehyde dehydrogenase 1 family, member L1 (*ALDH1L*) (**Figure 3C**). We performed DEX analysis between the two genotypes in each cell types that we identified. The majority of the DEX genes were found in the ExN clusters, with the number of DEX genes highest in ExN4 (**Figure 3 D**). Of the five cell types with the most DEX genes, four were excitatory neuron clusters (**Figure 3D,E and Supplemental Figure 3D**).

**Figure 3.**
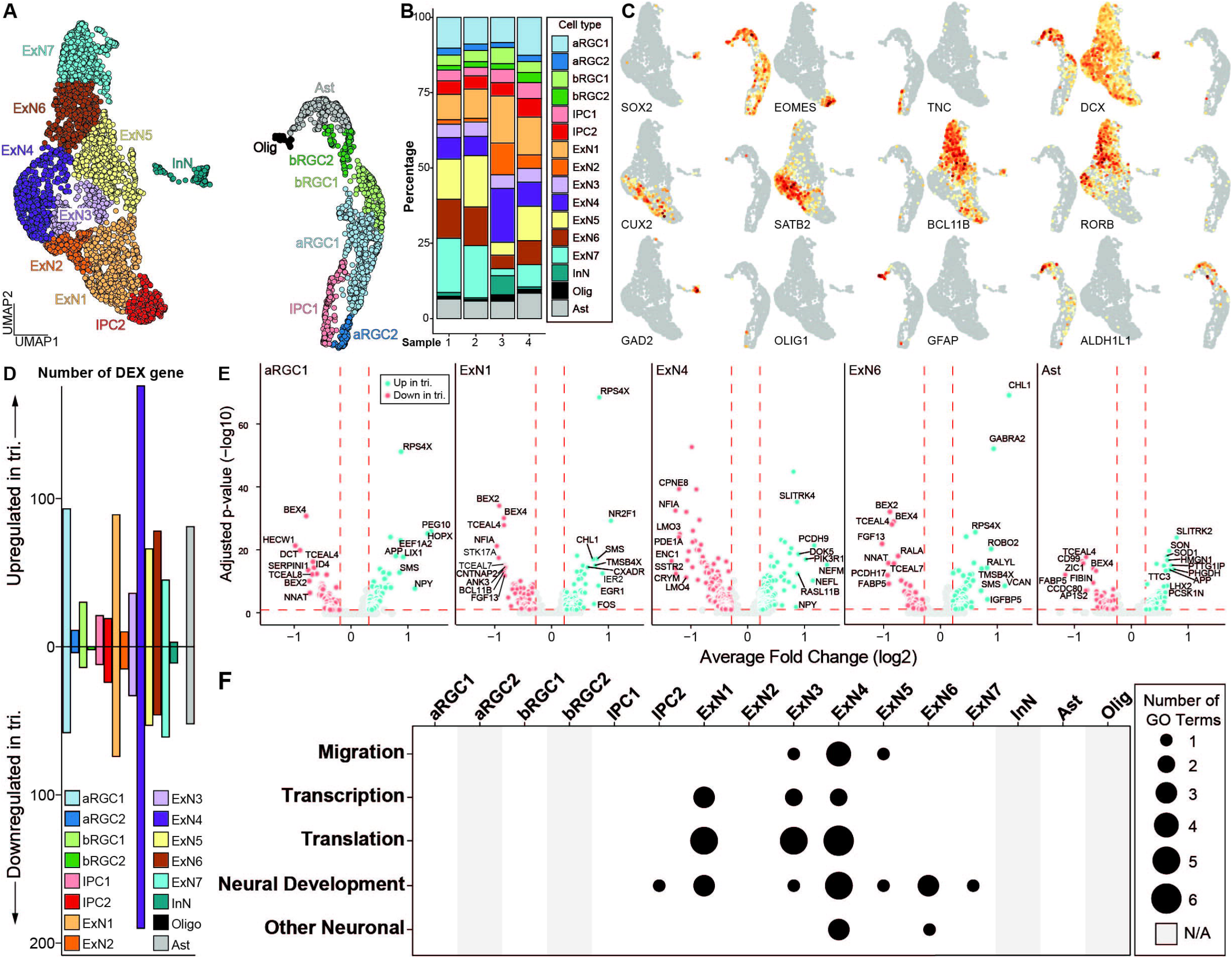
scRNA-seq analysis of isogenic euploid and trisomic CS at day 130. **(A)** UMAP representation of scRNA-seq data collected from two euploid and two trisomic CS samples. Colors represent identified cell types. ARGC, apical radial glial cell; bRGC, basal radial glial cell; IPC, intermedial progenitor cell; ExN, excitatory neuron; InN, inhibitory neuron; Ast, Astrocyte; Olig, oligodendrocyte. **(B)** Bar graph showing the percentage of each identified cell type in each sample. Colors represent same cell types as in (A). Sample 1 and 2 are euploid, whereas 3 and 4 are trisomic. **(C)** UMAP as in (A) showing gene expression levels of canonical markers. Colors represent normalized gene expression level (norm. exp.). **(D)** Bar graph showing the number of up- or down-regulated DEX genes in trisomic vs. euploid single cells by cell type. Colors represent same cell types as in (A). **(E)** Volcano plots showing DEX genes in cell types. Five cell types with the highest numbers of DEX genes are shown. Colors represent genotype (euploid, red; trisomic, blue). Vertical red dashed lines represent average log2 fold change of -0.25 or 0.25. Horizontal red dashed lines represent adjusted p-value of 0.1. Eu, euploid; tri, trisomy. **(F)** Dot plot showing the number of enriched gene ontology (GO) terms in each cell type. Enriched GO terms are grouped into five categories of “Migration”, “Transcription”, “Translation”, “Neural Development” and “Other Neuronal.” Size of the dot represents number of enriched GO terms. Grey bar represents cell types where no DEX genes were identified and thus not applicable (N/A) to the GO analysis.

We then performed gene ontology (GO) analysis to identify biological processes that are significantly enriched in each one of the cell types, using up and down regulated genes and a threshold of FDR < 0.05 to identify significantly affected biological processes (**Figure 3F and Supplemental Figure 4**). Thesebiological processes were further categorized into five groups: “migration”, “transcription”, “translation”, “other neuron related processes” and “neurodevelopment”. Consistent with the DEX analysis, excitatory neuron cell types showed the most significant enrichment with the highest number of enriched terms in all five categories, suggesting that DEX genes in ExN cell types converged on similar biological processes. In contrast, DEX genes from neural progenitor cell types (i.e. aRGC1, bRGC1 and IPC1) as well as Ast did not show any enrichment and thus had no functional convergence, even though the number of DEX genes was comparable to those from ExNs (**Figure 3D,F**).

We next performed pseudotime analysis to establish the differentiation trajectory for all cells (**Figure 4A and Supplemental Figure 5**). To quantify the difference driven by trisomy for each cell type, we calculated the average Euclidean distance on diffusion maps (DMs) between each trisomic cell and each euploid cell within the same cell type and used it as a presentation of transcriptome divergence between the genotypes (**Supplemental Figure 6**). We term this value “observed inter-genotype distance” or oIGD. To identify statistically significant oIGD, we randomized the genotype assignment 1000 times within each cell type and calculated a distribution of estimated IGD (eIGD). By comparing oIGDs to eIGDs, we identified seven cell types with statistically significant oIGDs indicating a significant transcriptomic divergence between the genotypes. These included – in the order of most significant to least significant - ExN4, ExN1, ExN6, ExN2, IPC2, ExN5, and ExN3 (**Figure 4B**). In contrast, other cell types (including ExN7) did not have significant oIGDs and thus did not show genotypic divergence in our dataset. We then further investigated ExN4, the cell type with the greatest genotypic divergence. In line with the IGD analysis, the cell density distribution of trisomic and euploid cells were significantly different on the DM and non-overlapping (**Figure 4C,E**). In contrast, trisomic and euploid cells from ExN7 completely overlapped in cell density distribution (**Figure 4D,F**). The cells from the two genotypes in ExN4 also show differences in pseudotime distribution, indicating a developmental asynchrony of the trisomic cells with their euploid counterparts (**Figure 4E**). In ExN7, however, the cells from the two genotypes are comparable in pseudotime distribution (**Figure 4F**). Together, this indicates that the development of some ExN clusters, including ExN4, is more significantly impacted by trisomy 21 than other cell types.

**Figure 4.**
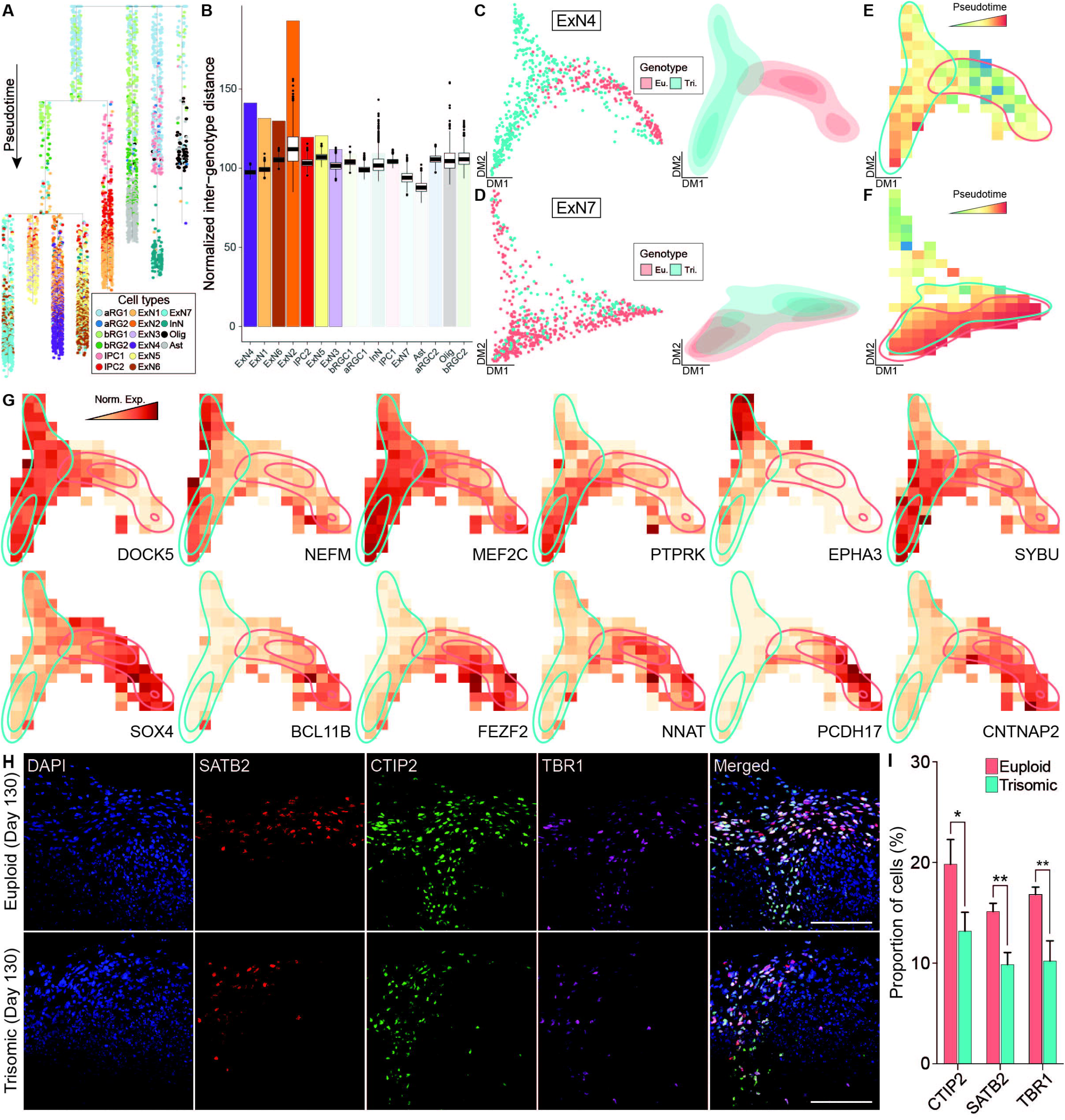
Pseudotime analysis of scRNA-seq data from euploid and trisomic CS at day 130. **(A)** Dendrogram showing single cells along pseudotime. Branches on dendrogram signify divergence in transcriptome profiles. Colors represent cell types. **(B)** Bar graph showing observed normalized inter-genotype distance (IGD) in each cell type. Box plots on top of each observed IGD show estimated IGDs from 1,000 permutations. Cell types are arranged by statistical significance of observed IGC. The first six cell types on the graph are statistically significant (p-value < 0.001). **(C)** Diffusion map (left panel) showing single cells and density plot (right) showing the distribution of single cells from ExN4 cell type. Colors represent genotype (eu., euploid, red; tri., trisomic, blue). **(D)** Diffusion map (left panel) showing single cells and density plot (right) showing the distribution of single cells from ExN7 cell type. Colors represent genotype (euploid, red; trisomic, blue). **(E)** Raster plot showing pseudotime in diffusion map space of ExN4 as in (C). Colors represent pseudotime. Regions with high density of euploid (red) or trisomic (blue) cells are outlined. **(F)** Raster plot showing pseudotime in diffusion map space of ExN7 as in (D). Heatmap colors represent pseudotime. Regions with high density of euploid (red) or trisomic (blue) cells are outlined. **(G)** Raster plot showing expression levels in diffusion map space as in (C) of genes specifically associated with trisomic or euploid cells in ExN4. Heatmap colors represent normalized gene expression levels (norm. exp.). Regions with high density of euploid (red) or trisomic (blue) cells are outlined. **(H)** IHC staining with anti-SATB2, andti-CTIP2 and anti-TBR1 antibodies in euploid and trisomic CS on day 130. **(I)** Bar graph showing the percentage of cells expressing CTIP2, SATB2 or TBR1 IHC signal over total number of cells stained with DAPI. The quantification is performed using ACEq application and analyzed using student t-test (euploid, n = 16; trisomic, n = 15). Error bar represents standard error. *p<0.05; **p<0.01. The quantification results are generated from three independent differentiation experiments; Scale bar, 100μm.

To identify genes driving the differences between the trisomic and euploid ExN4 cells, we performed principal graph analysis (PGA) (Cao et al., 2019). Superior to standard DEX analyses that are based solely on expression levels, PGA identifies genes not only by up- or downregulation between conditions, but also with nonrandom patterns along pseudotime, which we refer to as “association.” We identified ten genes that were specifically associated with trisomic cells in ExN4 including ephrin type-A receptor 3 (*EPHA3)* and myocyte enhancer factor 2C (*MEF2C*), which have been shown to function in motility and migration during neural development (**Figure 4G, Table 1 and Supplemental Figure 7)**. Among the genes unassociated with trisomic cells in ExN4 were several neuronal transcription factors such as *BCL11B* and *FEZF2*, as well as protocadherins, *PCDH17* and *PCDH19*, all of which play key roles in cortical development (**Figure 4G, Table 1 and Supplemental Figure 7**) (Chang et al., 2018; Du et al., 2021; Hoshina et al., 2021; Sadegh et al., 2021; Tsyporin et al., 2021). To validate the findings from scRNA-seq, we examined the protein expression of deep and superficial cortical layer markers, BCL11B (CTIP2) and SATB2, respectively in CS. Our analysis showed a significant decrease in the percentage of trisomic cells expressing CTIP2 (trisomic, 13.4±1.9%; euploid, 20±2.4%; p<0.04) as well as SATB2 (trisomic, 10%±1.1%; euploid, 15.34±0.8%; p<0.0015) at day 130 of differentiation. We also performed IHC staining for TBR1, a newborn neuron marker, and found 10.4±2% of trisomic cells were positive for the protein whereas 17±0.8% of euploid were positive (p<0.0008) at day 130 of differentiation **(Figure 4H,I**). The same reduction in the percentage of trisomic cells expressing these markers compared to euploid cells was observed on day 90 of differentiation: CTIP2 (trisomic, 21±4.4%; euploid, 39.4±3%; p<0.011), SATB2 (trisomic, 3.6±1.6%; euploid, 20.4±3.1%; p<0.0012), and TBR1 (trisomic, 17±5.2% euploid, 33.4±1.6%; p<0.032) **Supplemental Figure 1C,D)**. These data suggest that abnormal neurogenesis of excitatory neurons may also contribute to the reduction in trisomic CS volume, which is reminiscent of the reduction of cortical volume in individuals with trisomy 21.

**Table 1.** Genes from the principal graph analysis of ExN4 that are associated or unassociated with the trisomic genotype.

| Gene | Full name | Description | References |
| --- | --- | --- | --- |
| EPHA3 | EPH Receptor A3 | Receptor tyrosine kinase implicated in cell-cell adhesion, cell migration, and axon guidance | Yun et al., 2003; Brennaman et al., 2014 |
| DCLK3 | Doublecortin Like Kinase 3 | Predicted protein of the doublecortin superfamily | Mullins et al., 2021 |
| TGFB2 | Transforming Growth Factor Beta 2 | Secreted ligand of TGFβ proteins; Involved in SMAD signaling. | Wang et al., 2020 |
| NEFM | Neurofilament Medium Chain | Intermediate filament; plans a role in intracellular transport in axons and dendrites | Bergström et al., 2021 |
| PIK3R1 | Phosphoinositide-3-Kinase Regulatory Subunit 1 | Plays an important role in the metabolic actions of insulin | Jia et al., 2020 |
| DOK5 | Docking Protein 5 | Participates in RET-mediated neurite outgrowth | Liu et al., 2010 |
| PTPRK | Protein Tyrosine Phosphatase Receptor Type K | Regulates cell growth differentiation and mitosis | Shen et al., 1999 |
| MEF2C | Myocyte Enhancer Factor 2C | Transcription factor important for neocortical development | Li et al., 2018 |
| RPS4X | Ribosomal Protein S4 X-Linked | Ribosome involved in local translation in axon. | Shigeoka et al., 2019 |
| SYBU | Syntabulin | Component of kinesin motor complex for anterograde axonal transport | Berezcki et al., 2018 |
| SOX4 | SRY-Box Transcription Factor 4 | Transcription factor important for neurodevelopment | Shim et al., 2012; Da Silva et al., 2021 |
| PBX1 | PBX Homeobox 1 | Transcription factor implicated in regional patterning of the brain | Ypsilanti et al., 2021 |
| MEIS2 | Meis Homeobox 2 | Transcription factor essential for development | Matsuda et al., 2019 |
| VCAN | Versican | Proteoglycans of extracellular matrix | Armstrong et al., 2020 |
| IGSF3 | Immunoglobulin Superfamily Member 3 | Immunoglobulin-like membrane protein involved in neuronal morphogenesis | Usardi et al., 2017 |
| ENC1 | Ectodermal-Neural Cortex 1 | A member of the kelch family; interacts with actin | García-Calero and Puelles, 2005 |
| FEZF2 | FEZ Family Zinc Finger 2 | Transcription factor essential for projection neuron development | Sadegh et al., 2021; Tsyporin et al., 2021 |
| PCDH19 | Protocadherin 19 | A member of protocadherin subclass of the cadherin superfamily | Hoshina et al., 2021 |
| DUSP4 | Dual Specificity Phosphatase 4 | Mitogen Activated Protein Kinase (MAPK) inhibitor | Kirchner et al., 2020 |
| CNTNAP2 | Contactin Associated Protein 2 | Cell adhesion molecule of the neurexin family | Klibaite et al., 2022 |
| LMO7 | LIM Domain 7 | Protein-protein interaction | Lencer et al., 2017 |
| SSTR2 | Somatostatin Receptor 2 | Regulates neuronal calcium signalling | Agoglia et al., 2021 |
| PDE1A | Phosphodiesterase 1A | Important cellular signal transduction molecule | Stoner et al., 2014 |
| NFIA | Nuclear Factor I A | Transcription factor that regulates central nervous system development | Ogura et al., 2022 |
| BCL11B | BAF Chromatin Remodeling Complex Subunit BCL11B | Transcription factor regulating development of cortical projection neurons | Du et al., 2021 |
| NNAT | Neuronatin | Proteolipid involved in the regulation of ion channels | Kanno et al., 2019 |
| TOX | Thymocyte Selection Associated High Mobility Group Box | Transcription factor controlling proliferation of neural stem cells | Artegiani et al., 2017 |
| CRYM | Crystallin Mu | Binds to thyroid hormone and regulates neurodevelopment by binding to thyroid hormone | Hallen and Cooper, 2017 |
| PCDH17 | Protocadherin 17 | A member of protocadherin subclass of the cadherin superfamily important for establishing cell-cell connections in the brain | Chang et al., 2018 |
| CPNE8 | Copine 8 | Calcium-dependent membrane-binding protein | Florentinus-Mefailoski et al., 2021 |

### 3.4 Comparison with other transcriptomic studies reveals common and diverged gene dysregulation in trisomy 21 CS

To put our findings in a broader context, we compared our current scRNA-seq dataset to three previously published DS transcriptomics datasets, including bulk RNA-seq data of brain-like NPCs differentiated from iPSCs (Klein et al., 2022), bulk microarray data of postmortem human brain (Olmos-Serrano et al., 2016), and scRNA-seq data of postmortem brains (Palmer et al., 2021). It is worth noting that Klein et al., 2022 dataset was generated from the same isogenic line (WC-24-02-DS-B (euploid) and WC-24-02-DS-M (trisomic)) we used to generate the CS in our current study. In order to make the comparison to the Klein et al., 2022 dataset more meaningful, only data from iPSCs 8 days after induction (representing neural progenitor cells) were used. In the Palmer et al., 2021 study, we analyzed both the full dataset which consisted of samples from multiple age groups and both sexes (“all”), as well as a selection of the older female samples (“old fem.”) to examine the potential confounds from mixing different biological sexes. We first investigated the DEX genes from each study located on chromosome 21 (HSA21) and found statistically significant overlaps between every pair of datasets with *p* < 0.001 (**Figure 5A,B**). Particularly, 45 (88%) and 43 (84%) out of the 51 HSA21 DEX genes from our current dataset were also DEX in the Klein et al., 2022 and Palmer et al., 2021 (old fem.) datasets, respectively. In addition, 18 (35%) of 51 the HSA21 DEX genes from our dataset were also DEX in the Olmos-Serrano et al., 2016 dataset. Remarkably, we found 16 DEX genes on HSA21 that were present in all four datasets (**Figure 5C and Table 2)**.

**Figure 5.**
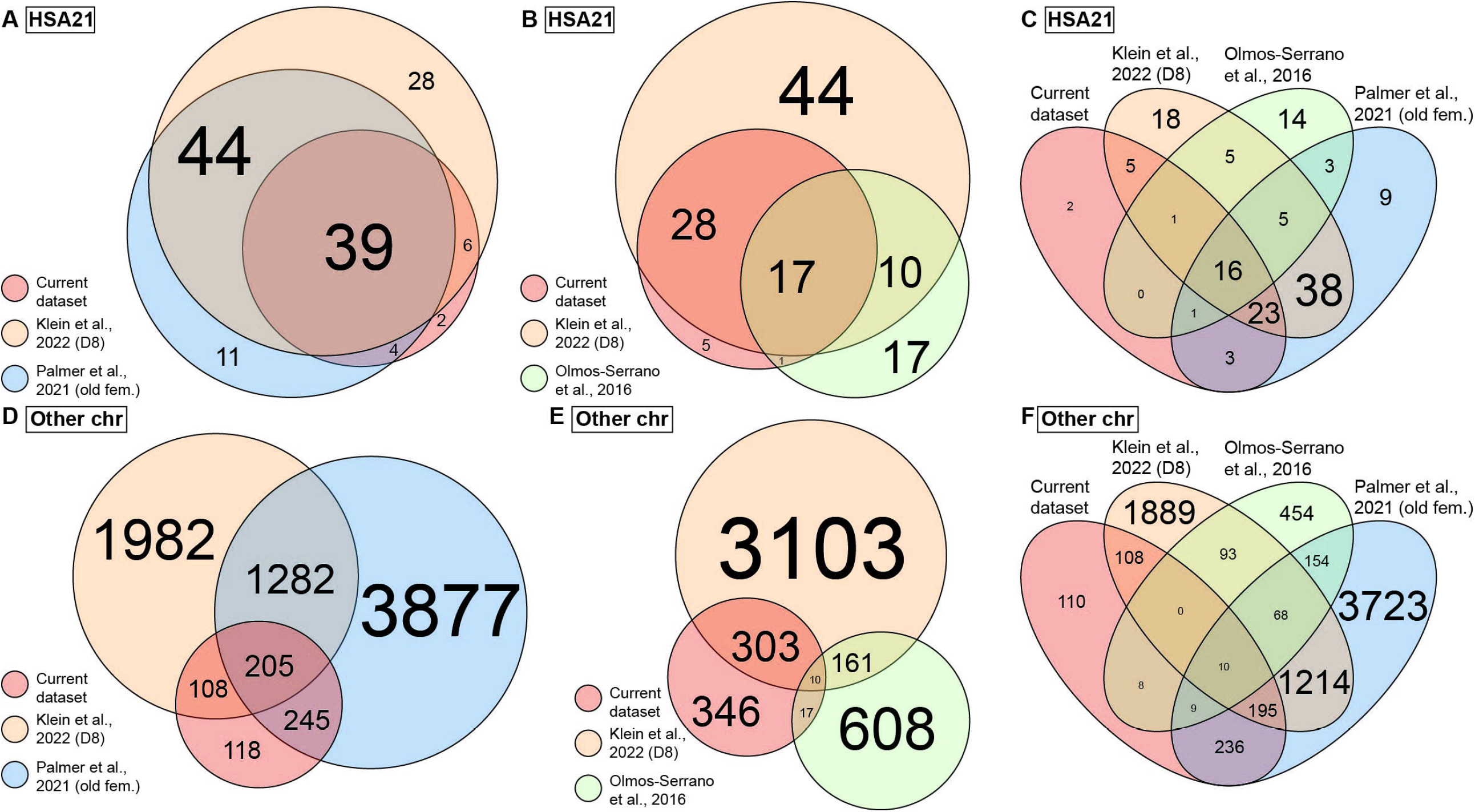
Overlap of DEX genes between the current and published datasets. **(A)** Venn diagram showing overlap of DEX genes on chromosome 21 (HAS21) between the current, Klein et al., 2022 and Palmer et al., 2021 datasets. Circle size represents number of DEX genes. **(B)** Venn diagram showing overlap of DEX genes on HAS21 between the current, Klein et al., 2022 and Olmos-Serrano et al., 2016 datasets. Circle size represents number of DEX genes. **(C)** Venn diagram showing overlap of DEX genes on HAS21 between all four datasets as in (A) and (B). **(D)** Venn diagram showing overlap of DEX genes on all chromosomes except HAS21 between the current, Klein et al., 2022 and Palmer et al., 2021 datasets. Circle size represents number of DEX genes. **(E)** Venn diagram showing overlap of DEX genes on all chromosomes except HAS21 between the current, Klein et al., 2022 and Olmos-Serrano et al., 2016 datasets. Circle size represents number of DEX genes. **(F)** Venn diagram showing overlap of DEX genes on all chromosomes except HAS21 between all four datasets as in (D) and (E). Colors represent datasets. Only data of old female (old fem.) samples are included from Palmer et al., 2021 dataset and only data of day 8 WC-24-02-DS (D8) iPSC cultures are included from Klein et al., 2022. Size of the text in all panels represents number of DEX genes.

**Table 2.** Sixteen DEX genes on HSA21 that were present in all four datasets.

| Gene | Full name | Description | Reference |
| --- | --- | --- | --- |
| SOD1 | Superoxide Dismutase | A cellular antioxidant, breaks down reactive oxygen species | Rosen et al., 1993; Milani et al., 2011 |
| PCNT | Pericentrin | Plays a role in the organization of the centromeres and mitotic spindle formation. | Chen et al., 1996 |
| PFKL | Phosphofructokinase | Participates in glucose metabolism | Levanon et al., 1989 |
| GART | Trifunctional purine biosynthetic protein adenosine-3 | Important in the biogenesis of purines | Gnirke et al., 1991 |
| PRMT2 | Protein Arginine Methyltransferase 2 | Plays an important role in the metabolism and formation of nuclear pre-mRNA | Scott et al., 1998 |
| PDXK | Pyridoxal Kinase | Plays a role in vitamin B6 metabolism | Hanna et al., 1997 |
| SON | SON DNA and RNA Binding Protein | Splicing co-factor contributing to efficient splicing of cell cycle regulators | Ahn et al., 2011 |
| ITSN1 | Intersectin 1 | Regulates endocytic trafficking and actin polymerization | Guipponi et al., 1998 |
| CRYZL1 | Crystallin Zeta Like 1 | Regulates glucose metabolism and lipogenesis. | Kim et al., 1999 |
| BRWD1 | Bromodomain And WD Repeat Domain Containing 1 | Participates in a formation of multicomplex proteins and participates in epigenetic regulation | Huang et al., 2003; Mandal et al., 2018 |
| TMEM50B | Transmembrane Protein 50B | Predicted to regulate late endosome and multivesicular body formation and sorting. | Moldrich et al., 2008; Kong et al., 2014 |
| MRPS6 | Mitochondrial Ribosomal Protein S6 | Participates in the protein synthesis within the mitochondrion | Oviya et al., 2021 |
| PttG1IP | Pituitary Tumor-Transforming Gene 1 Protein-Interacting Protein | Interacts with a proto-oncogene, PTTG1 and plays a role in cancer | Chien and Pei, 2000 |
| IFNAR1 | Interferon Alpha And Beta Receptor Subunit 1 | A part of the interferon pathway. Upon activation stimulates Janus protein kinase | Novick et al., 1994; Shemesh et al., 2021 |
| IFNAR2 | Interferon Alpha And Beta Receptor Subunit 2 | A part of the interferon pathway. Upon activation stimulates Janus protein kinase and controls STAT phosphorylation | Shemesh et al., 2021 |
| USP16 | Ubiquitin Specific Peptidase 16 | Deubiquitinating enzyme participating in the metaphase to anaphase transition in mitosis | Puente et al., 2003 |

**Table 3.**
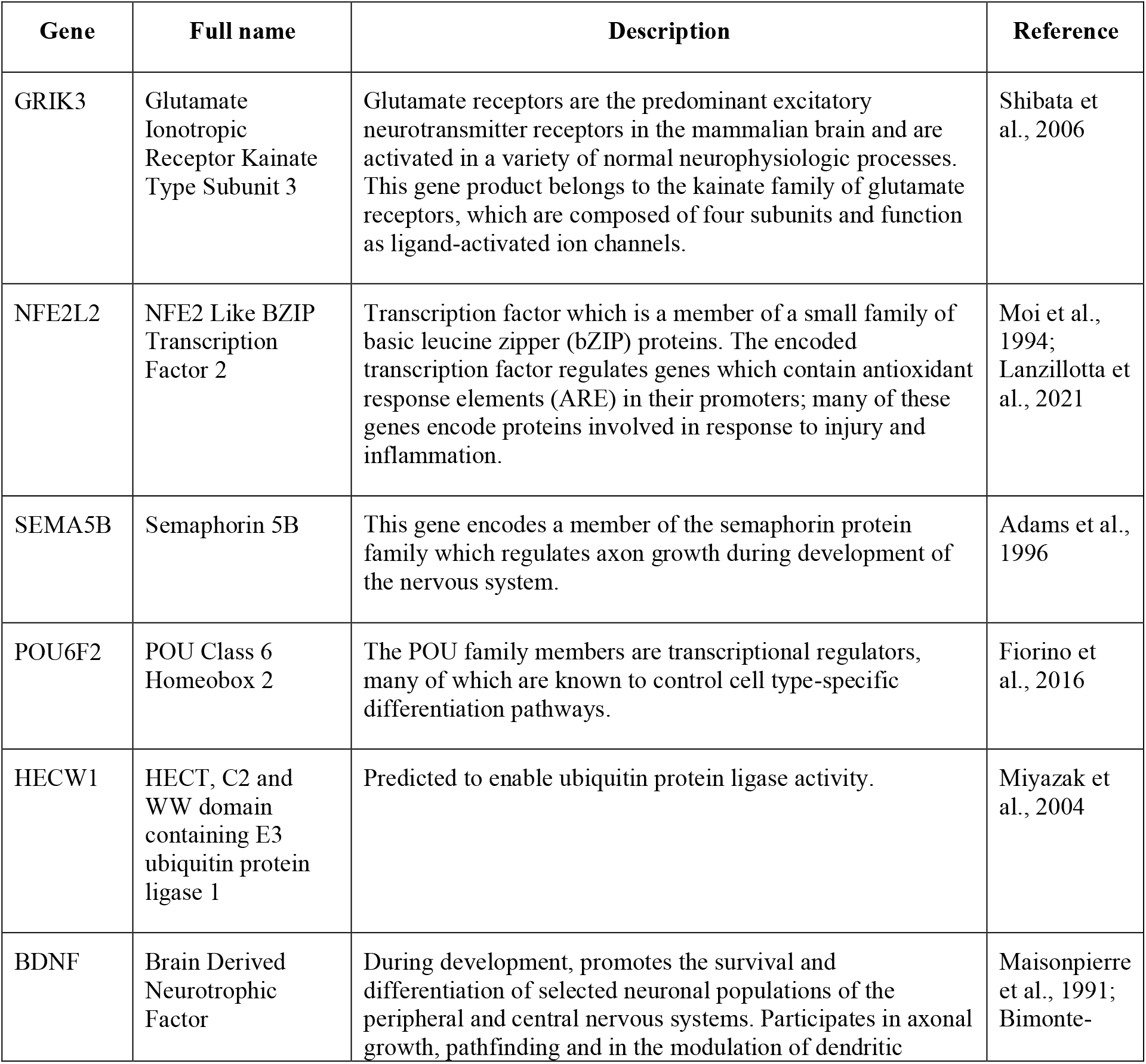
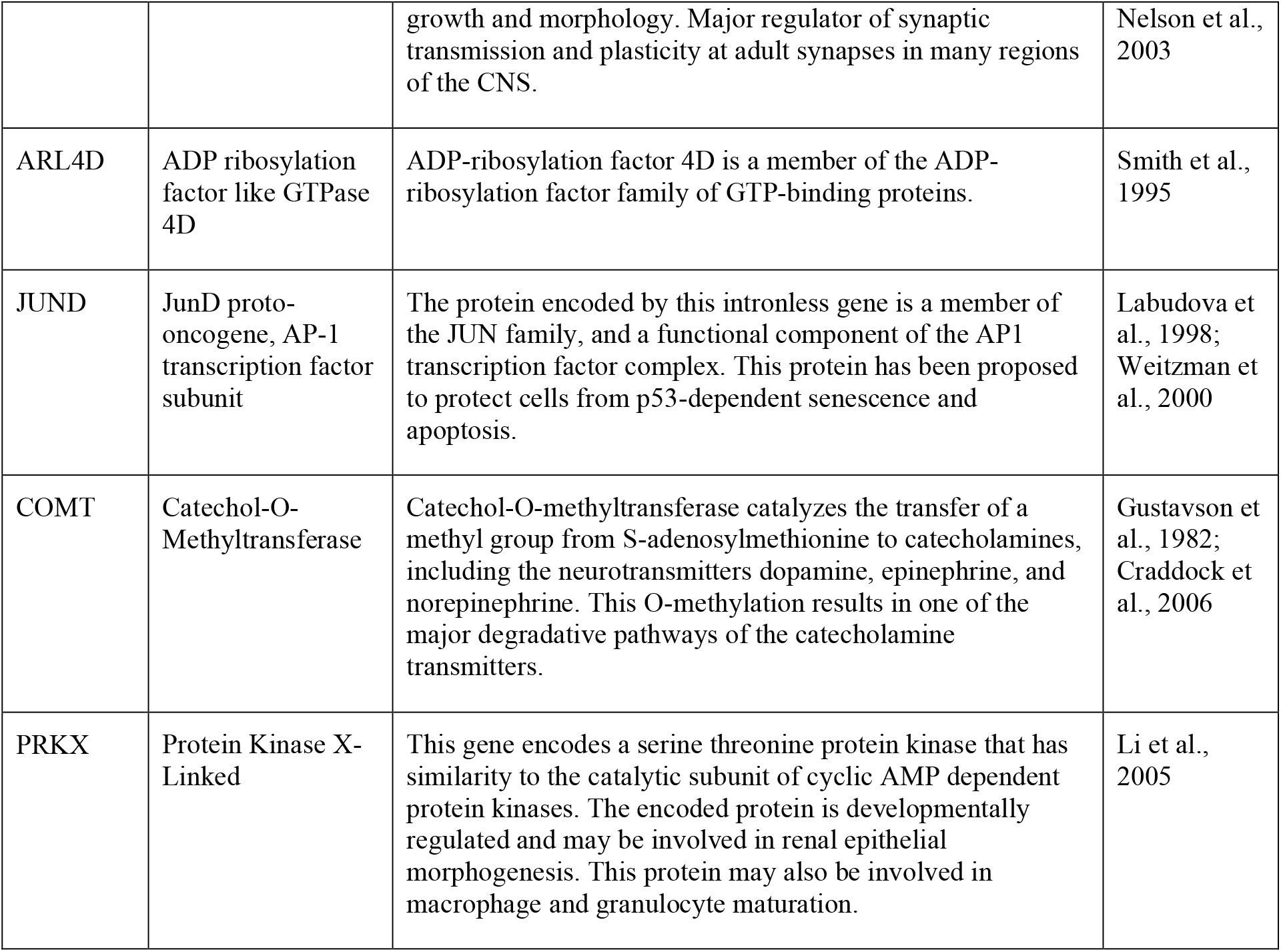
Genes that are DEX in common across all four datasets: Klein et al. 2022, Palmer et al. 2021, Olmos-Serrano et al. 2016, and the current scRNA-seq dataset generated for this study.

We next examined the DEX genes on chromosomes other than HSA21 (non-HSA21). Here, 46% and 67% of the DEX genes from our dataset overlapped with those identified in the Klein et al., 2022 and Palmer et al., 2021 datasets (old fem.), respectively, and both overlaps were statistically significant (*p* < 0.001; **Figure 5D**). About 4% of the DEX genes from our dataset overlapped with the Olmos-Serrano et al., 2016 dataset, which was not statistically significant (*p* = 0.449; **Figure 5E**). Despite the technical and biological differences of the datasets, we were able to identify 10 non-HSA21 genes that were DEX in all four datasets, some of which have previously been linked to DS or other neurodevelopmental phenotypes in DS (**Table 2**). To assess the influence of variability in individual genome on DEX gene discovery, we repeated the analyses replacing Olmos-Serrano et al., 2016 dataset with Palmer et al., 2021 (all) dataset (**Supplemental Figure 8A-D**). We first observed that Palmer et al., 2021 contained fewer unique HSA21 DEX genes (16 of 111, 14.4%) than Olmos-Serrano et al. 2016 (17 of 45, 37.8%), when compared to our dataset and Klein et al., 2022 (**Supplemental Figure 8A**). As expected, most HSA21 DEX genes (95 of 98, 96.9%) from Palmer et al., 2021 (old fem.) were also DEX in Palmer et al., 2021 (all) (**Supplemental Figure 8B**). Interestingly, when we compared non-HAS21 DEX genes from Palmer et al., 2021 (all) to other datasets, we found substantially more unique DEX genes. In fact, 6031 (71.2%) non-HSA21 DEX genes were unique to Palmer et al., 2021 (all) when compared to our dataset and Klein et al., 2022 dataset (**Supplemental Figure 8C**). In addition, 5429 (64.1%) non-HSA21 DEX genes were also unique in Palmer et al., 2021 (all) when Palmer et al., 2021 (old fem.) dataset was included (**Supplemental Figure 8D**). These observations suggest that multiple sample sources and variability in individual genomes contribute significantly to transcriptomic differences.

## 4 Discussion

In this study, we generated CS from isogenic iPSC lines derived from an adult female with DS to examine changes in neurodevelopment at cellular resolution. The CS we generated as a model of dorsal forebrain development recapitulate the environment of fetal human neocortex at mid-gestation (Qian et al., 2019), including major cell types such as RGC, IPC, ExN, InN and glial cells. This allows for a detailed examination of aberrant neurodevelopmental processes not only at a cellular level but also without the confound of multiple genetic backgrounds in a mixed sample set. Importantly, our trisomic CS captured one of the most salient features of neurodevelopment in DS that is thought to underlie the development of the intellectual disability in DS, a decrease in volume, which is reminiscent of the decreased cortical volume in brains of individuals with DS (Wisniewski, 1990; Golden and Hyman, 1994; Pinter et al., 2001; McCann et al., 2021). While we identified increased apoptosis as one of the contributing factors to the decreased volume, we hypothesized that changes in neuronal differentiation and cortical expansion, another well-defined phenotype in DS, also contributed to the smaller size of the CS (Schmidt-Sidor et al., 1990; Guidi et al., 2008; Larsen et al., 2008).

Both histological and scRNA-seq analyses point specifically to changes in excitatory neuron development in the trisomic CS. IHC analysis identifies a significant decrease in percentage of TBR1+ neurons at day 90 and 130 indicating that there are fewer newly differentiated neurons over an extended period during which neurons are born *in vitro*. There were also significant decreases in BCL11B (CTIP2) expression, a marker of mature neocortical layer V pyramidal neurons, and SATB2 a marker of layer II-IV neocortical excitatory neurons at the time points we tested. The decrease in neuronal marker gene and protein expressions indicate that cell populations in trisomic CS resembling both upper and deep layer neurons in the brain appear to be affected, supporting previous fundings in DS-derived organoids (Tang et al., 2021). Using IGD, we identified transcriptional divergence between genotypes within multiple neuronal cell types and found that ExN cell types in general were most severely affected by HSA21 trisomy. In fact, six of the seven cell types with statistically significant IGD are ExN, consistent with the observation that four out of five cell types with the most DEX genes were ExN. Additionally, 47 of the 48 significantly enriched GO terms identified by our analyses were enriched in ExN cell types.

Although changes in neurogenesis have been described previously, the underlying molecular mechanism of these changes is unknown (Tang et al., 2021). In our study, we identified ExN4 as the most severely affected ExN cell type, as it had the largest number of DEX genes and the most significant inter-genotype distance among all cell types. Therefore, we focused our analysis on ExN4 to further understand molecular changes due to HSA21 in a cell type specific manner. Firstly, ExN4 expressed *SATB2* and *RORB4*, the combination of which indicate that these cells share transcriptomic signatures with layer IV excitatory neurons in human neocortex. The fact that ExN4 is the most profoundly affected cell type in our analysis may suggest that layer IV neocortical excitatory neurons in individuals with DS are also under higher pathological risk than other cell types during fetal development, supporting previous findings (Tang et al., 2021). Secondly, pseudotime analysis also indicated that there were changes in developmental trajectory of the trisomic cells. This developmental asynchrony between cells of the two genotypes in ExN4 may be the result of transcriptional dysregulation leading to abnormal maturation. Indeed, based on our PGA results, several genes were associated strongly with the trisomic ExN4 cells while other genes were strongly unassociated with trisomic ExN4 cells. For instance, *BCL11B* and *Family Zinc Finger 2* (*FEZF2*), which are important transcription factors during fetal brain development, are unassociated with trisomic ExN4 cells. Both BCL11B and FEZF2, while commonly recognized as markers for layer V pyramidal neurons in postnatal neocortex, have transient broader expression patterns during fetal development in immature neurons (Du et al., 2021). During development, BCL11B is essential for the formation and maintenance of synapses (Simon et al., 2012) and its deficiency is associated with intellectual deficits, developmental delay and speech impairment (Punwani et al., 2016; Lessel et al., 2018). FEZ2F is expressed in postmitotic neurons and it regulates the acquisition of cell identity and specification in cortical projection neurons through the repression of alternative neuronal fate genes (Chen et al., 2005; Molyneaux et al., 2005; Shim et al., 2012; Tsyporin et al., 2021). Therefore, our observation that *BCL11B* and *FEZF2* are strongly unassociated with trisomic ExN4 cells may indicate a premature consolidation and shutdown of necessary transcriptional programs due to HSA21 trisomy. The premature shutdown of transcriptional program may in turn lead to alterations in the development of individual neurons, which may then manifest either in programed cell death as we observed earlier, or in changes in neural plasticity and connectivity as suggested by previous human (Suetsugu and Mehraein, 1980; Ferrer and Gullotta, 1990; Medalla et al., 2017) and mouse (Dierssen et al., 2003; Belichenko et al., 2004; Pérez-Cremades et al., 2010) studies.

Lastly, among the genes strongly associated with the trisomic genotype in ExN4 are *EPHA3*, a gene critical for cytoskeleton organization, migration and cell adhesion of neural cells during nervous system development (Rudolph et al., 2010; Javier-Torrent et al., 2019) and *MEF2C*, an important transcription factor regulating early neuronal differentiation and cortical lamination (Li et al., 2008). Interestingly, *EPHA3* was also identified as a DEX gene in trisomic excitatory neurons in a recent study (Palmer et al., 2021). It is noteworthy that Palmer et al. performed scRNA-seq on postmortem brain tissue from individuals with DS, which is vastly different from the *in vitro* samples we used in the current study. However, despite the biological differences, both studies identified *EPHA3* as a downstream factor of HSA21 trisomy, suggesting that cell-cell signaling, and neuronal motility may be a common aspect of DS pathology. It also supports that the technical and quantitative approaches that the two studies employed were rigorous and replicable, and that CS as a model system for brain development is valid and promising.

Along the same line, we observed great overlap of DEX genes between our study and published studies, once again providing evidence for the validity of our experimental system and approach. Furthermore, by focusing our study on isogenic cell lines, we eliminated confounding biological factors such as sex, ethnicity, somatic variability, etc. To demonstrate this unique aspect of our study, we compared the current dataset to three previously published transcriptomics datasets. We first examined DEX genes on HSA21 and found statistically significant overlap between every pair of datasets, including 16 genes that were shared by all datasets. The observation suggests that a largely consistent cohort of HSA21 genes are dysregulated in neocortex or cortical model systems, regardless of age, sex or experimental system. Since dysregulation of HSA21 is the root cause of DS phenotypes, the remarkable overlap of HSA21 DEX genes we observed not only cross-validates the findings of the included studies, but also highlights the 16 shared DEX genes as key candidates for DS in the brain. Conversely, the fact that out of all triplicated HSA21 genes only 16 were present in all 3 datasets implies that gene dosage effect in trisomy 21 varies between cell types, tissue, and developmental stage.

Comparing non-HSA21 DEX genes from the studies, we once again identified statistically significant overlap between DEX genes from our CS dataset and those from the Klein et al. 2022 (D8), as well as those from Palmer et al., 2021 (all and old fem.). We also found significant overlap between the Palmer et al., 2021 dataset with the Klein et al., 2022 dataset. The only dataset that did not have statistically significant overlap with our current was Olmos-Serrano et al. 2016. Similarly, the overlap between the Klein et al., 2022 dataset with Olmos-Serrano et al. 2016 was also much weaker. Since Olmos-Serrano et al. 2016 was the only dataset generated with microarray technology, the lack of overlap in non-HSA21 DEX genes may be largely due to technical differences between microarray and RNA-seq approaches, especially considering that both the Olmos-Serrano et al. 2016 and Palmer et al. 2021 (all) dataset are similarly composed of samples from multiple individuals across different ages and of both sexes.

Next, we wanted to understand the impact of biological sex on transcriptional variability in the studies. In the current study, we focused our analysis on isogenic lines of CS derived from a female individual. By doing so, we eliminated the influence of sex on the transcriptome, which allowed us to distill transcriptomic changes specific to the woman the sample was derived from. To illustrate the influence of sex on transcriptomics data, we reanalyzed scRNA-seq data of only the female samples from the Palmer et al., 2021 (age range of 39 to 60) and compared it to the entire Palmer et al. 2021 dataset with samples from both sexes. Surprisingly, we observed that while 5429 genes were DEX in the Palmer et al. 2021 (all), 3280 genes were DEX in Palmer et al. 2021 (old fem.), and only 2143 DEX genes were shared between the two. These findings support the effect biological sex and age has on the transcriptome of individuals with DS, demonstrating that these variables should be controlled in future studies, regardless of whether they are focused on gene expression or on cellular/anatomical datasets.

Another aspect of experimental design that may influence the power of the analyses is the variability of individual genetic background of the subjects. We sought to minimize the effect of individual genetic background by studying isogenic cell lines derived from the same individual. We demonstrated the advantage of this approach by comparing DEX genes we identified in the current study to those identified in Klein et al., 2022, in which the same isogenic lines were used. While the cell lines were exposed to completely different culturing and differentiation conditions and were analyzed by different technologies, we observed the most significant overlap of DEX genes between our study and Klein et al., 2022. Furthermore, even though we observed nearly perfect overlap of DEX genes on HSA21, the overlap for non-HSA21 genes (while still significant), was less substantial. This indicates that features of dysregulation across the genome are affected by background genetics and environmental conditions. Therefore, an attention to genetic predisposition and addressing the individualized aspects of DS in a “personalized” manner should be an important consideration for future studies.

Despite the differences between datasets, we identified ten genes on non-HSA21 chromosomes that were dysregulated all studies: *GRIK3, NFE2L2, SEMA5B, POU6F2, HECW1, BDNF, ARL4D, JUND, COMT*, and *PRKX*. As these ten genes are consistently dysregulated across age, sex, sample type, and sequencing technology, they may strongly impact the consistent neurodevelopmental phenotypes in DS leading to the intellectual disability.

A few of these genes have been reported before in conjunction with DS. Increased levels of *COMT*, encoding catechol-O-methyltransferase, has been reported in erythrocytes in individuals with DS (Gustavson et al., 1982). Intriguingly, this enzyme is active in the prefrontal cortex and is responsible for metabolizing catecholamine neurotransmitters. Mutations in *COMT* have previously been associated with executive dysfunction and schizophrenia (Bearden et al., 2004; Baker et al., 2005; Craddock et al., 2006). Increased levels of COMT activity in DS may disrupt homeostatic levels of important neurotransmitters, impairing neural transmission. NFE2L2 has previously been implicated in the development of Alzheimer’s pathology in DS (Sharma et al., 2020; Lanzillotta et al., 2021) and decreased mRNA and protein levels of BDNF have long been linked to deficits in learning and memory in mouse models of DS (Bimonte-Nelson et al., 2003; Bianchi et al., 2010; Parrini et al., 2017). Decreased levels of JUND have been reported in samples from brains of individuals with DS (Labudova et al., 1998). As JUND has been shown to protect cells from apoptosis (Weitzman et al., 2000), decrease in its expression is in line with the increase in apoptosis we observed in our trisomic CS.

The other six DEX genes in common from all four datasets have not been previously associated with DS. However, most of them have been shown to play important roles in neurodevelopment and their dysregulation in DS may be a common mechanism contributing to the neurological changes and ID in all individuals with DS. *POU6F2* encodes a transcription factor important in neural subtype determination (Fiorino et al., 2016; Ainsworth et al., 2018). *GRIK3* encodes a glutamate receptor subunit. Gain of function mutations of *GRIK3* have previously been associated with ID and neurodevelopmental deficits (Shibata et al., 2006). *SEMA5B* is important for axon guidance and cell migration, which we also identified as key dysregulated features in ExN from our trisomic CS (Yazdani and Terman, 2006). The recurrent dysregulation of the ten DEX genes across multiple datasets suggest their function and dysfunction may be key to understanding common aspects of the neurodevelopmental deficits across all individuals with the neurodevelopmental deficits characteristic of DS.

Altogether our study demonstrates the power of deeply analyzing genetically defined isogenic CS in a personalized manner, in conjunction with broad examination of transcriptional dysregulation, in the context of DS. As transcriptional dysregulation in neurodevelopmental diseases such as DS varies not only between individuals but also between tissue and cell types, it is also important to examine transcriptomic changes at the cellular level to gain functionally relevant and clinically actionable insights. At the same time, the variability across technological and experimental conditions and its influence on the interpretation of results should not be underestimated. Our study identifies multiple genes of interest consistent across datasets. Future studies will be necessary to confirm and elucidate their role in neurodevelopmental phenotypes in DS.

## 5 Data Availability Statement

The scRNA-seq data for this study can be found in the Sequence Read Archive (SRA) with access number PRJNA828127.

## 7 Author Contributions

E.Z. and T.F.H conceived the idea. J.A.K., E.Z. and R.K. performed all the tissue culture experiments. The bioinformatics analysis was done by Z.L., S.R., and Y.P assisted with data analysis. Z.L., J.A.K., E.Z., and T.F.H wrote the manuscript.

## 9 Acknowledgments

We would like to acknowledge Dr. Anita Bhattacharyya of the University of Wisconsin for her kind gift of the WC-24-02-DS-B and WC-24-02-DS-M isogenic iPSCs. We thank the sequencing core at Boston University School of Medicine, Boston University for their help in library preparation and sequencing. We also thank Neuroinformatics Workgroup at the Center for Neuroscience Research, Children’s National Research Institute, for the transcriptomic analysis.

## 10 Permission to reuse and Copyright

Figure 1 was created with BioRender.com. Jenny Klein has been granted license number VW23RFOF5H to permits BioRender content to be sublicensed for use in journal publications.

## Footnotes

1 https://www.bumc.bu.edu/anatneuro/ella-zeldich-lab/

