## Supplementary material for "Asynchronous excitatory neuron development in an isogenic cortical spheroid model of Down syndrome": Li et al., 2022, Supplementary material: Li_et_al_2022_Supplementary_Material.pdf

### Supplementary Figures

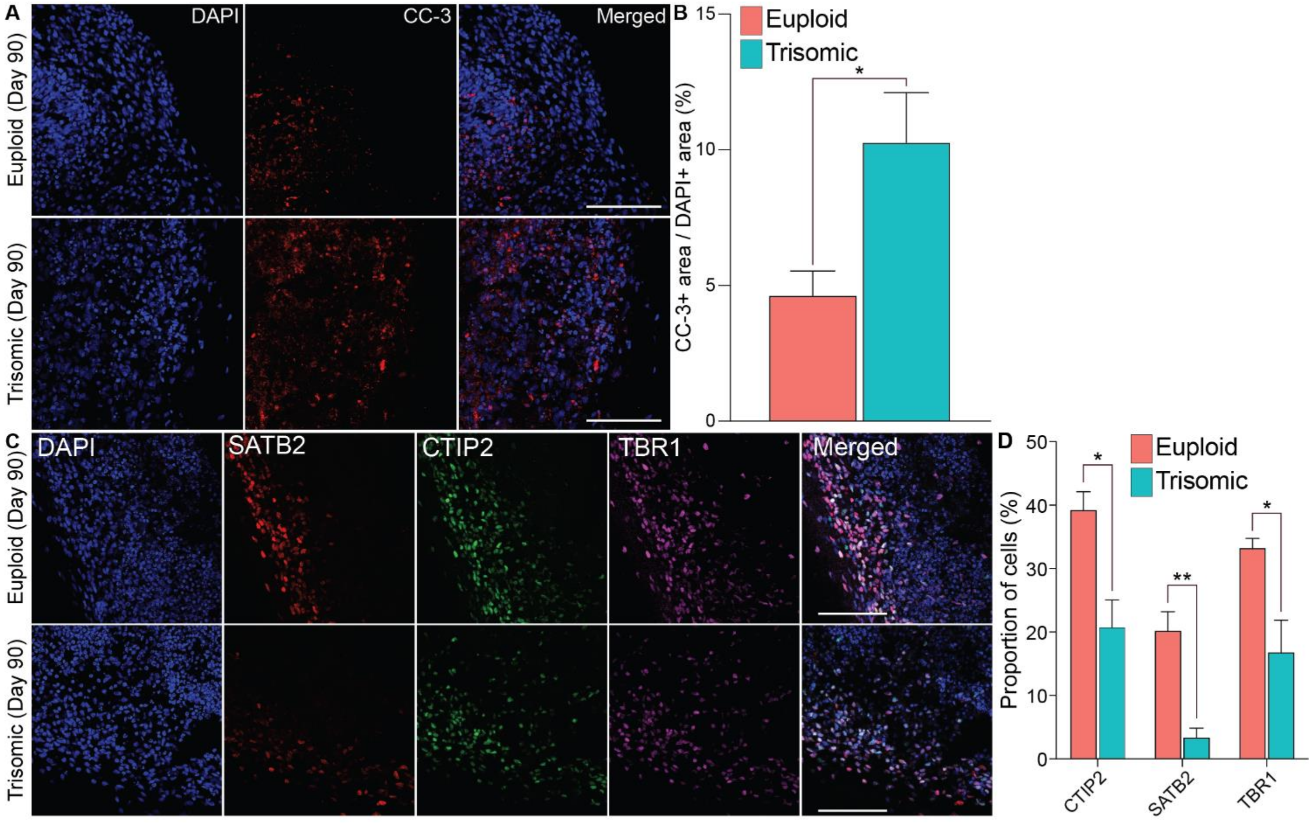


**Supplementary Figure 1. Cell death and neurogenesis in euploid and trisomic CS on day 90. (A)** IHC staining of euploid and trisomic CS stained with the anti-CC-3 and anti-Tuj1 antibodies on day 90. **(B)** Bar graph showing the percentage of area with CC-3 IHC signal over the area with DAPI signal, quantified through particles analysis via ImageJ and analyzed using student t-test; (euploid, n = 12; trisomic, n = 12). **(C)** IHC staining of euploid and trisomic CS anti-SATB2, anti-CTIP2, and anti-TBR1 antibodies on day 90. **(D)** Bar graph showing the percentage of cells expressing CTIP2, SATB2 or TBR1 IHC signal over total number of cells stained with DAPI. The quantification is performed using ACEq application and analyzed using student t-test (euploid, n = 12; trisomic, n = 12). Error bars represent standard error. *p<0.05, **p<0.01, ***p<0.001. The quantification results are generated from three independent differentiation experiments. Scale bar, 100µm.

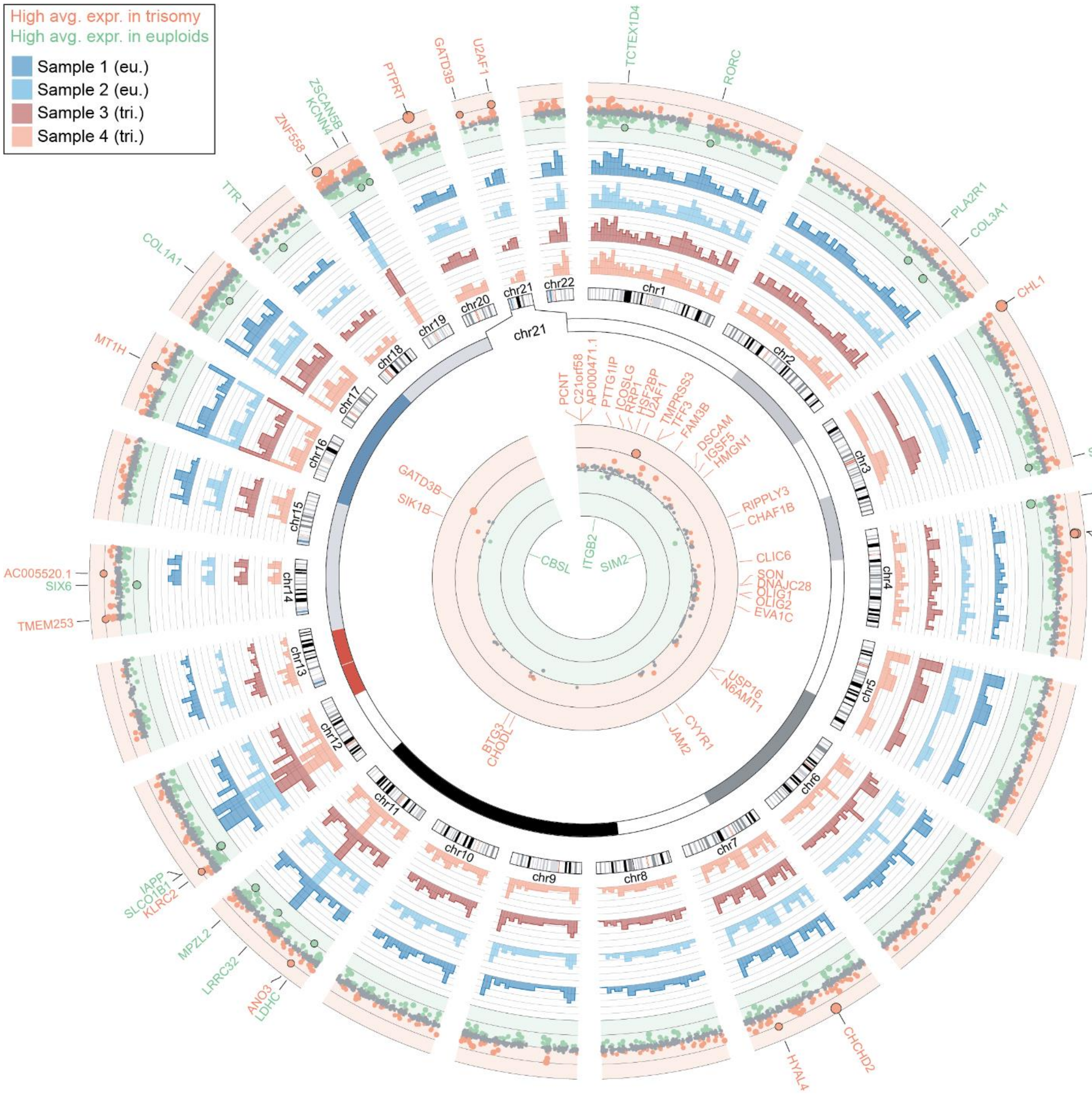


**Supplementary Figure 2. Coverage of reads over autosomes from scRNA-seq analysis of euploid and trisomic cortical spheroid CS on day 130.** Circos plot showing properties of scRNA-seq data from euploid and trisomic CS. The outermost track shows genes with higher average level of expression (avg. expr.) in trisomic (red) or euploid (green) samples as scatter plots. Dots further away from the mid-line and greater in size represent genes with greater fold change. Dots representing genes with fold change greater than two are colored. Those with fold change greater than 55 are outlined in black and are labeled with gene name. The next four tracks show read coverage over autosomes in each sample in five million base pair windows as bar plots. The height of the bar represents level of raw reads counts. Euploid samples (eu.) are colored in blue and trisomic samples (tri.) in red. Diagrams of autosomes are shown next to the bar plots. The innermost track is a zoom-in of chromosome 21 (chr21). Genes with higher avg. expr. in trisomic (red) or euploid (green) samples are shown as scatter plots. Genes with fold change greater than 2 are colored and labeled.

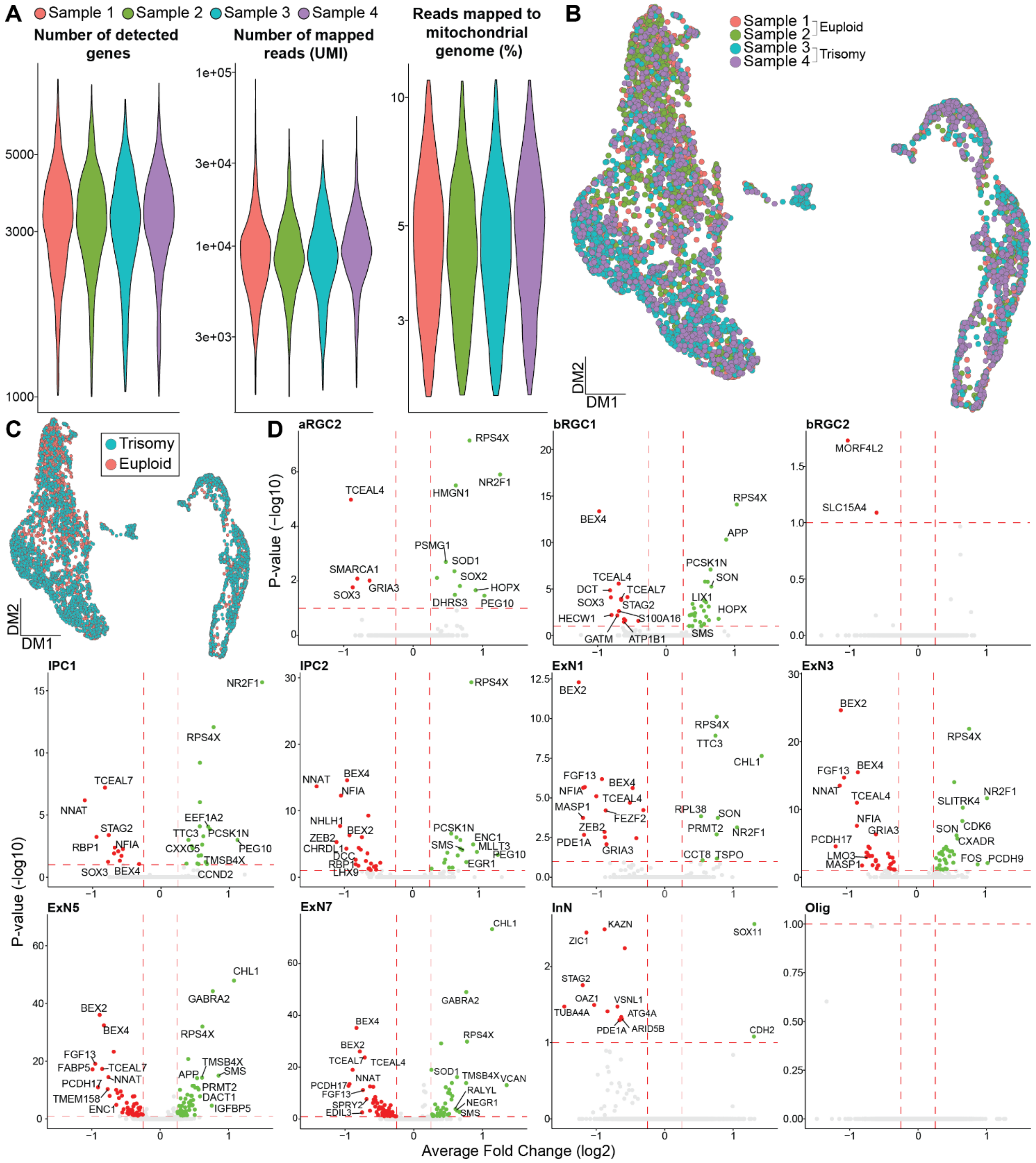


**Supplementary Figure 3. Quality control of scRNA-seq datasets of euploid and trisomic cortical spheroid on day 130. (A)** Violin plots showing number of detected genes, number of mapped reads (UMI) and percentage of reads mapped to mitochondrial genome in each sample. Colors represent samples. **(B)** UMAP of the scRNA-seq dataset colored by samples. **(C)** UMAP of the scRNA-seq dataset as in (B) colored by genotype (euploid, red; trisomic, blue). **(D)** Volcano plots showing DEX genes in cell types not included in Figure 3E. Colors represent genotype (euploid, red; trisomic, blue). Vertical red dashed lines represent average log2 fold change of -0.25 or 0.25. Horizontal red dashed lines represent adjusted p-value of 0.1.

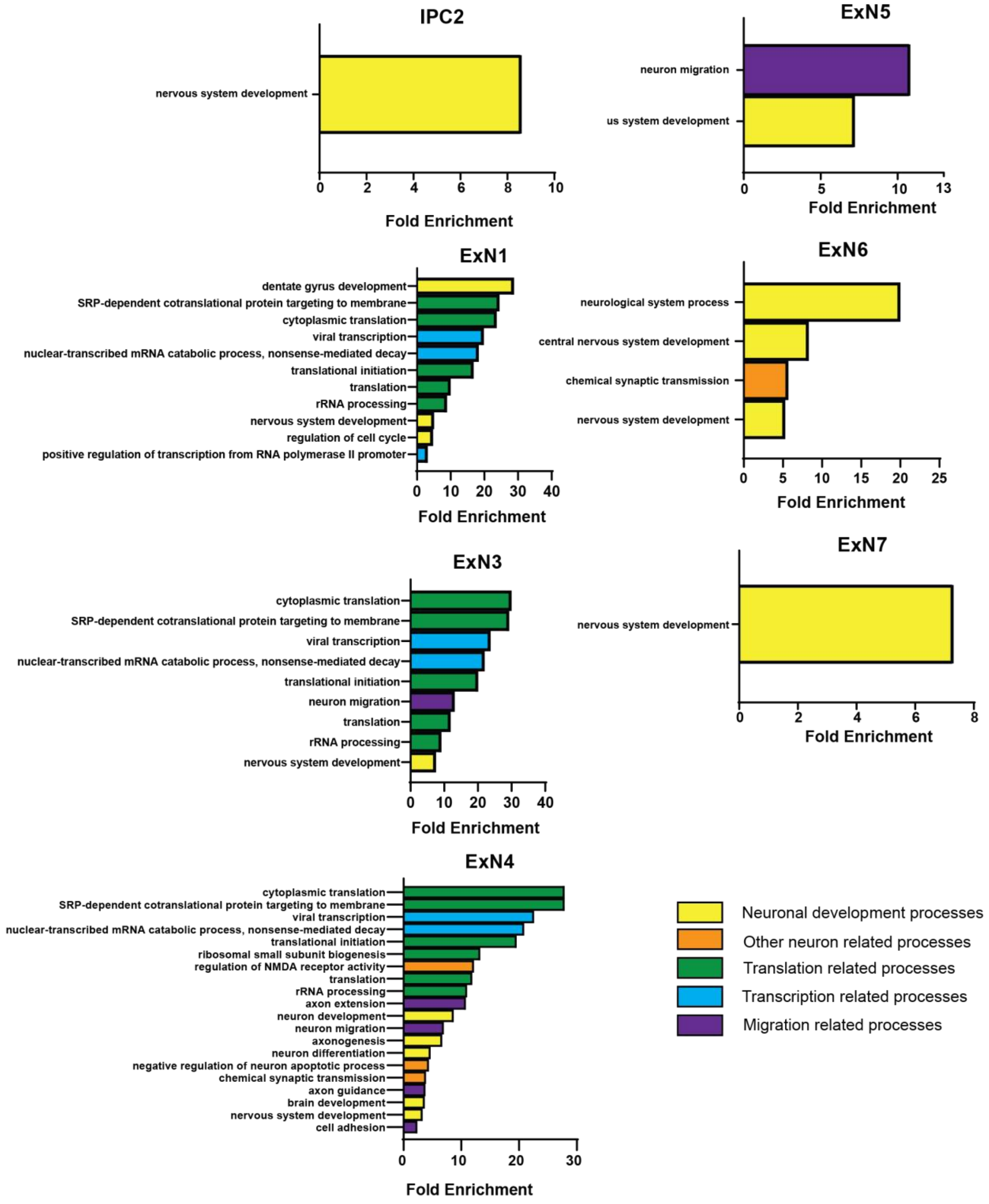


**Supplementary Figure 4. GO analysis of DEX genes in cell types identified scRNA-seq analysis of euploid and trisomic CS samples on day 130.** Bar plots showing significant fold enrichment of GO terms in each cell type based on DEX genes comparing trisomic to euploid CS scRNA-seq data. Only cell types with significantly enriched GO terms are shown. Colors represent categories of GO terms.

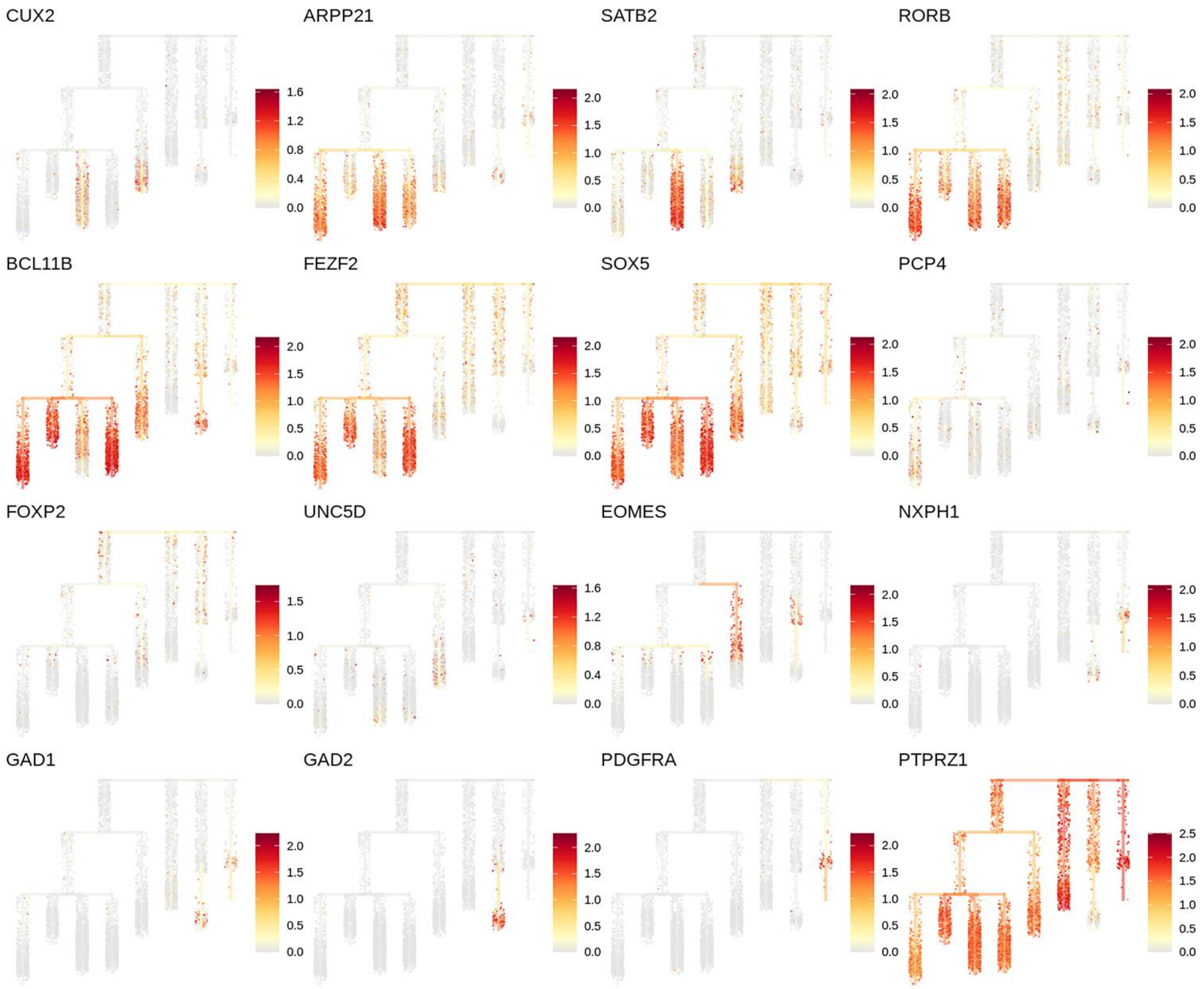


**Supplementary Figure 5. Canonical marker gene expression along pseudotime in euploid and trisomic CS scRNA-seq data on day 130.** Dendrogram showing single cells along pseudotime. Branches on dendrogram signify divergence in transcriptome profiles. Heatmap colors represent levels of normalized gene expression.

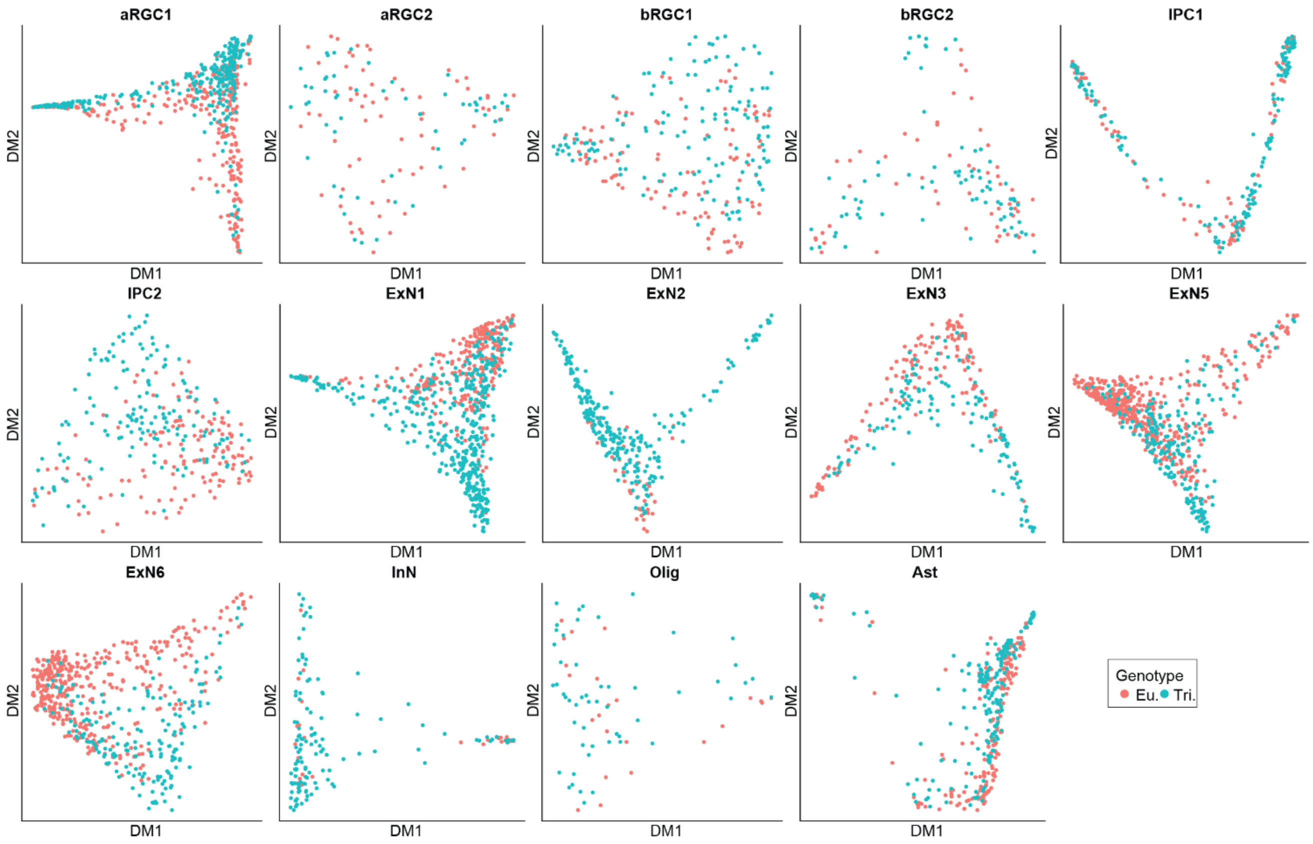


**Supplementary Figure 6. Diffusion maps depicting scRNA-seq data from euploid and trisomic CS on day 130 by cell type.** Diffusion maps for cell types not shown in Figure 4C and D are presented. Colors represent genotype (euploid, red; trisomic, blue).

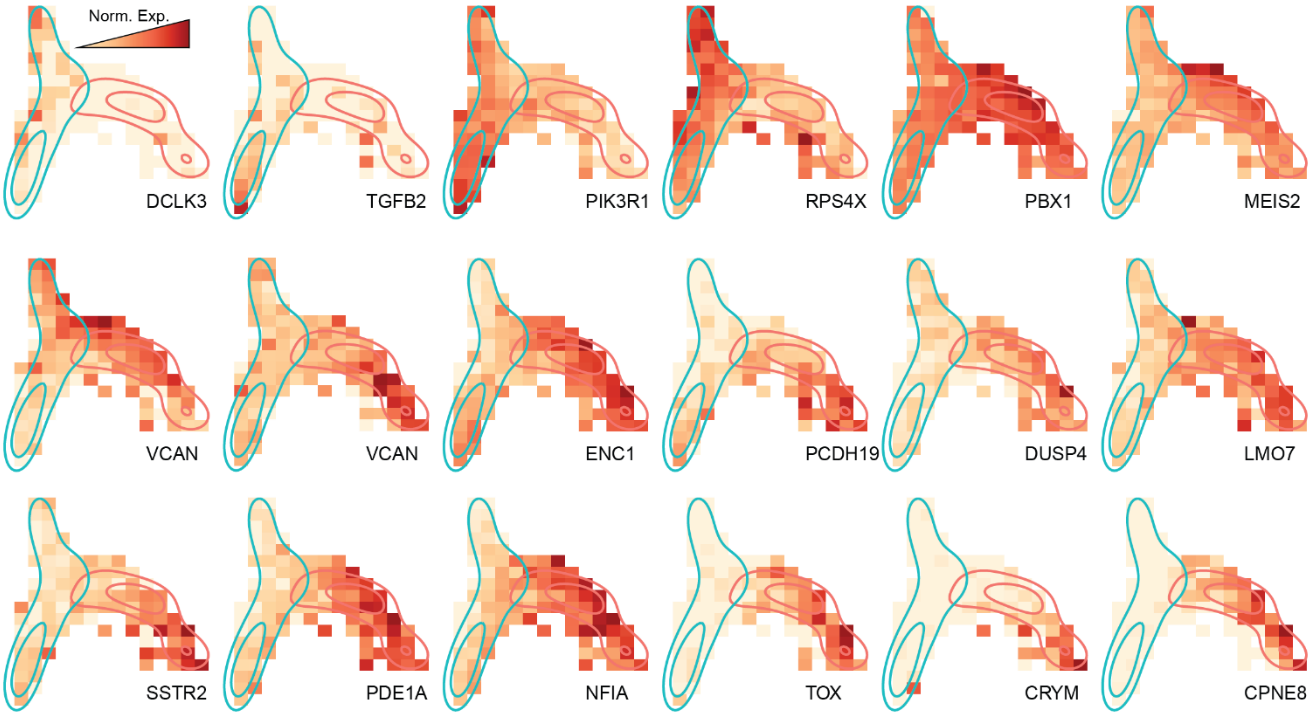


**Supplementary Figure 7. Genotype-driven differences along pseudotime in scRNA-seq data from euploid and trisomic CS on day 130.** Raster plot showing expression levels in diffusion map space as in (C) of genes specifically associated with trisomic or euploid cells in ExN4. Heatmap colors represent normalized gene expression levels (norm. exp.). Regions with high density of euploid (red) or trisomic (blue) cells are outlined.

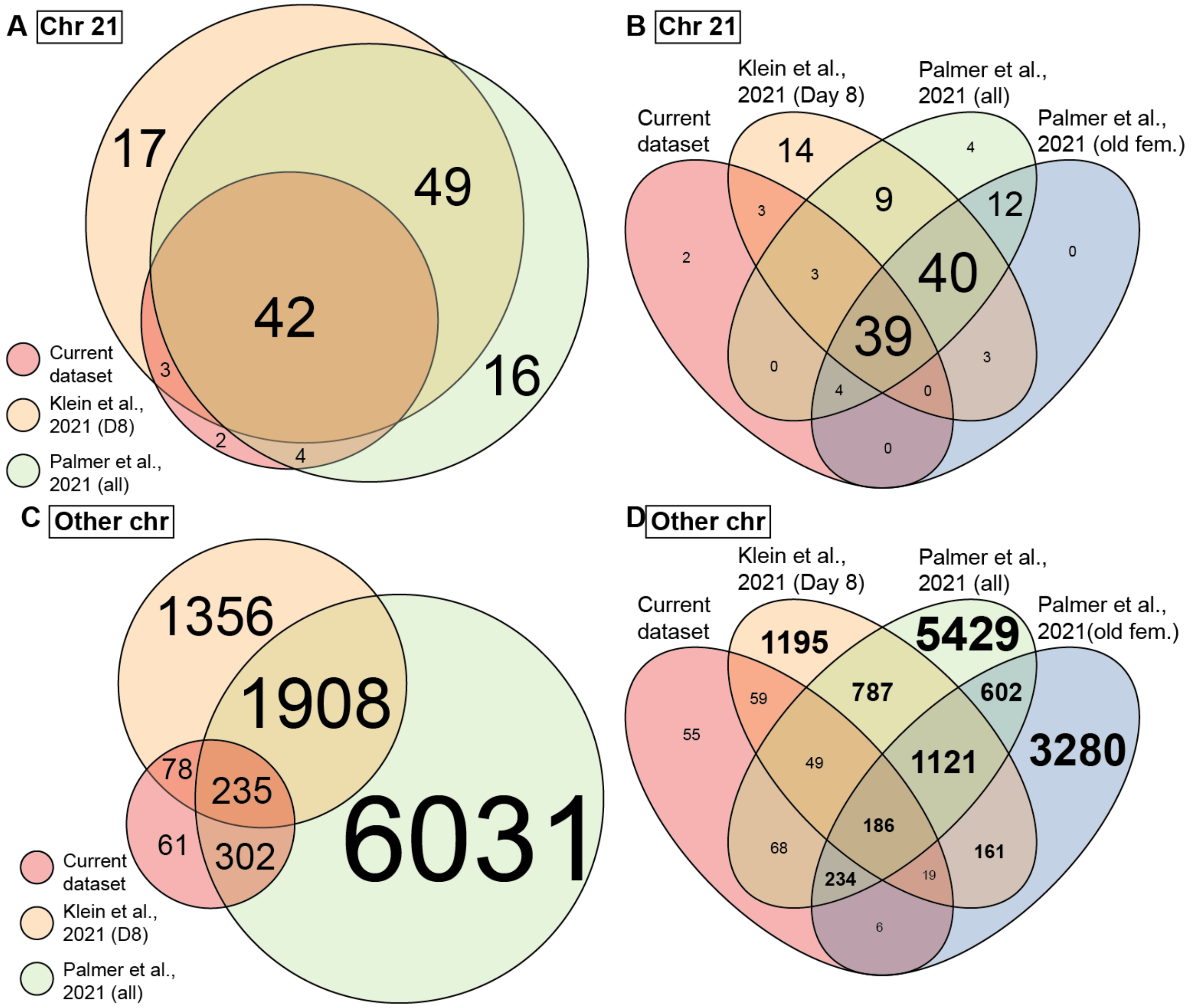


**Supplementary Figure 8. Overlap of DEX genes between the current and published datasets. (A)** Venn diagram showing overlap of DEX genes on HAS21 between the current, Klein et al., 2022 (D8) datasets and Palmer et al., 2021 datasets. Circle size represents number of DEX genes. Only data of old female (old fem.) samples are included from Palmer et al., 2021 dataset and only data of day 8 WC-24-02-DS (D8) iPSCs cultures are included from Klein et al., 2022. All samples from Palmer et al., 2021 (all) are included **(B)** Venn diagram showing overlap of DEX genes on HAS21 between four datasets as in (A) and (B). Only data of old female samples from Palmer et al., 2021 (all) dataset are included in Palmer et al., 2021 (old fem.). **(C)** Venn diagram showing overlap of DEX genes on all chromosomes except HAS21 between the current, Klein et al., 2022 (D8) datasets and Palmer et al., 2021 (all) datasets. Size of circle represents number of DEX genes. **(D)** Venn diagram showing overlap of DEX genes on all chromosomes except HAS21 between all four datasets. Colors represent datasets. Size of the text in all panels represents number of DEX genes.
